# Integrative metagenomics and metabolomics reveal age-associated gut microbiota and metabolite alterations in experimental COVID-19

**DOI:** 10.1101/2024.11.05.622058

**Authors:** Patrícia Brito Rodrigues, Vinícius de Rezende Rodovalho, Valentin Sencio, Nicolas Benech, Marybeth Creskey, Fabiola Silva Angulo, Lou Delval, Cyril Robil, Philippe Gosset, Arnaud Machelart, Joel Haas, Amandine Descat, Jean François Goosens, Delphine Beury, Florence Maurier, David Hot, Isabelle Wolowczuk, Harry Sokol, Xu Zhang, Marco Aurélio Ramirez Vinolo, François Trottein

## Abstract

Aging is a key contributor of morbidity and mortality during acute viral pneumonia. The potential role of age-associated dysbiosis on disease outcomes is still elusive. In the current study, we used high-resolution shotgun metagenomics and targeted metabolomics to characterize SARS-CoV-2-associated changes in the gut microbiota from young (2-month-old) and aged (22-month-old) hamsters, a valuable model of COVID-19. We show that age-related dysfunctions in the gut microbiota are linked to disease severity and long-term sequelae in older hamsters. Our data also reveal age-specific changes in the composition and metabolic activity of the gut microbiota during both the acute phase (day 7 post-infection, D7) and the recovery phase (D22) of infection. Aged hamsters exhibited the most notable shifts in gut microbiota composition and plasma metabolic profiles. Through an integrative analysis of metagenomics, metabolomics, and clinical data, we identified significant associations between bacterial taxa, metabolites and disease markers in the aged group. On D7 (high viral load and lung epithelial damage) and D22 (body weight loss and fibrosis), numerous amino acids, amino acid-related molecules, and indole derivatives were found to correlate with disease markers. In particular, a persistent decrease in phenylalanine, tryptophan, glutamic acid, and indoleacetic acid in aged animals positively correlated with poor recovery of body weight and/or lung fibrosis by D22. In younger hamsters, several bacterial taxa (*Eubacterium*, *Oscillospiraceae*, *Lawsonibacter*) and plasma metabolites (carnosine and cis-aconitic acid) were associated with mild disease outcomes. These findings support the need for age-specific microbiome-targeting strategies to more effectively manage acute viral pneumonia and long-term disease outcomes.

## Introduction

Older adults (over ∼65 years) are particularly susceptible to respiratory viral infections, including those caused by influenza A viruses and severe acute respiratory syndrome coronavirus 2 (SARS-CoV-2), the etiologic agent of coronavirus disease-19 (COVID-19)^1–3^. This greater susceptibility to viral infections in the elderly is related to declines in pulmonary function and immune response, as well as altered recovery, prolonging the duration of sequelae^4–6^. It is well established that the gut microbiota plays a key role in the lung’s defense against respiratory viruses^7,8^. Recent clinical studies have also shown that alterations of the gut microbiota’s functionality during influenza and COVID-19 correlate with disease severity and/or long-term sequelae^9–24^. However, human studies investigating the effects of viral pneumonia on the pre-existing altered microbiota in older adults are sparse. We herein investigated for the first time the effects of a respiratory virus (SARS-CoV-2) on the gut microbiota’s functionality in a preclinical model of advanced aging. To this end, the Syrian golden hamster, a valuable model of COVID-19 research^25,26^, was used.

Aging associates with perturbations in gut microbiota composition and functionality^27,28^. In both humans and mice, aging is associated with a decline in microbial diversity and richness, including a notable decrease in beneficial bacteria like the short-chain fatty acid (SCFA)-producing *Lachnospiraceae*, *Ruminococcaceae*, and *Bifidobacteriaceae* family members^28–33^. Conversely, there is an increased relative abundance of the Pseudomonadota phylum (formerly termed as Proteobacteria), which may drive inflammaging^28^. Importantly, aging is associated with defects in multiple crucial metabolic functions of the gut microbiome. Several metabolic pathways are affected including those involved in the production of SCFAs, tryptophan metabolites (indole), and secondary biliary acid^28–33^. Interestingly, some products of these metabolic pathways play a role in the control of respiratory viral infections in both mice and humans^34–39^. Despite these clear alterations of the aged gut microbiota, few studies have explored the link between age-related dysbiosis and susceptibility (and severity) to viral respiratory infections. Fuentes and colleagues reported that the abundances of specific bacteria, including butyrate-producers and members of *Gammaproteobacteria* family, could serve as potential biomarkers for susceptibility to influenza infection in aged^15^. In a clinical study on COVID-19, MacCann and colleagues showed that the proliferation of opportunistic pathogens, such as *Clostridium hathewayi*, *Enterococcus faecium*, *Coprobacillus, Eggerthella*, *Actinomyces spp*., *Ruminococcus gnavus*, *Ruminococcus torques*, and *Bacteroides dorei*, correlated with dysregulated inflammatory response and disease severity in aged individuals^20^. SARS-CoV-2 has been shown to alter the gut microbiota composition in hamsters^26,40,41^, although the impact of advanced age remains unknown. In the current study, we investigated the relationship between age, gut microbiota, and SARS-CoV-2 infection, focusing on: (1) the association between age-related changes in gut microbiota and COVID-19 susceptibility and outcomes, and (2) the differential impact of SARS-CoV-2 infection on the gut microbiota of young adults versus aged individuals, and its potential influence on disease parameters. To investigate this, two-time points were studied: one corresponding to the peak of disease severity and another related to disease resolution. We combined shotgun metagenomics, targeted metabolomics, and measurements of COVID19 severity. This multi-omics approach allowed us to identify a specific signature potentially linked to gut microbiota that is associated with disease outcomes in advanced aged individuals.

## Results

### Aged hamsters exhibit alterations of gut microbiota composition and functionality at steady state

The aging process affects the composition and functional activity of the gut microbiota in both humans and rodents^42–44^ but its impact on hamster’s gut microbiota has not been studied until now. To address this knowledge gap, cecal samples were collected from young (2-month-old) and aged (22-month-old) hamsters and processed for metagenomics analysis. Shotgun metagenomic sequencing showed no significant differences in bacterial alpha diversity between the two age groups (**Supplementary Figure 1a**). However, beta diversity analysis, using an Aitchison distance-based principal coordinate analysis (PCoA), revealed significant differences in the bacterial communities between young and aged hamsters (*P* = 0.013, permutational multivariate analysis of variance, PERMANOVA) (**Figure 1a**). Consistent with previous studies using 16S rRNA gene sequencing^26,40,41^, shotgun analysis revealed that Bacillota (formerly Firmicutes) and Bacteroidota (Bacteroidetes) dominated the gut microbiota in hamsters. There was no significant change in their abundance with aging (73.3/69.6% and 24.5/27.5% in young/aged hamsters, respectively) (**Figure 1b**). At the genus level, *UBA9475*, *Oscillibacter*, *Alistipes*, *Acetatifactor*, *Amulumruptor*, and *Bacteroides* were the most abundant taxa, with no significant differences between the young and aged groups (**Supplementary Figure 1b**). A linear regression model revealed significant differences in the abundance of specific bacterial species between young and aged hamsters (**Figure 1c** and **Supplementary Table 1**). The main change at the order level was an increase in *Lactobacillales* abundance in aged hamsters. Compared to young animals, aged hamsters had higher relative abundances of *Bifidobacterium_animalis*, *Ligilactobacillus_murinus, Ligilactobacillus*_MGBC161554, *Parasutterella*_sp900552195, and *Alistipes sp.* (**Figure 1c**). In the young group, enriched species included several *Ruminiclostridium* strains and members of the *Lachnospiraceae* family (*e.g.,* species CAG-127_sp002493625).

**Figure 1.**
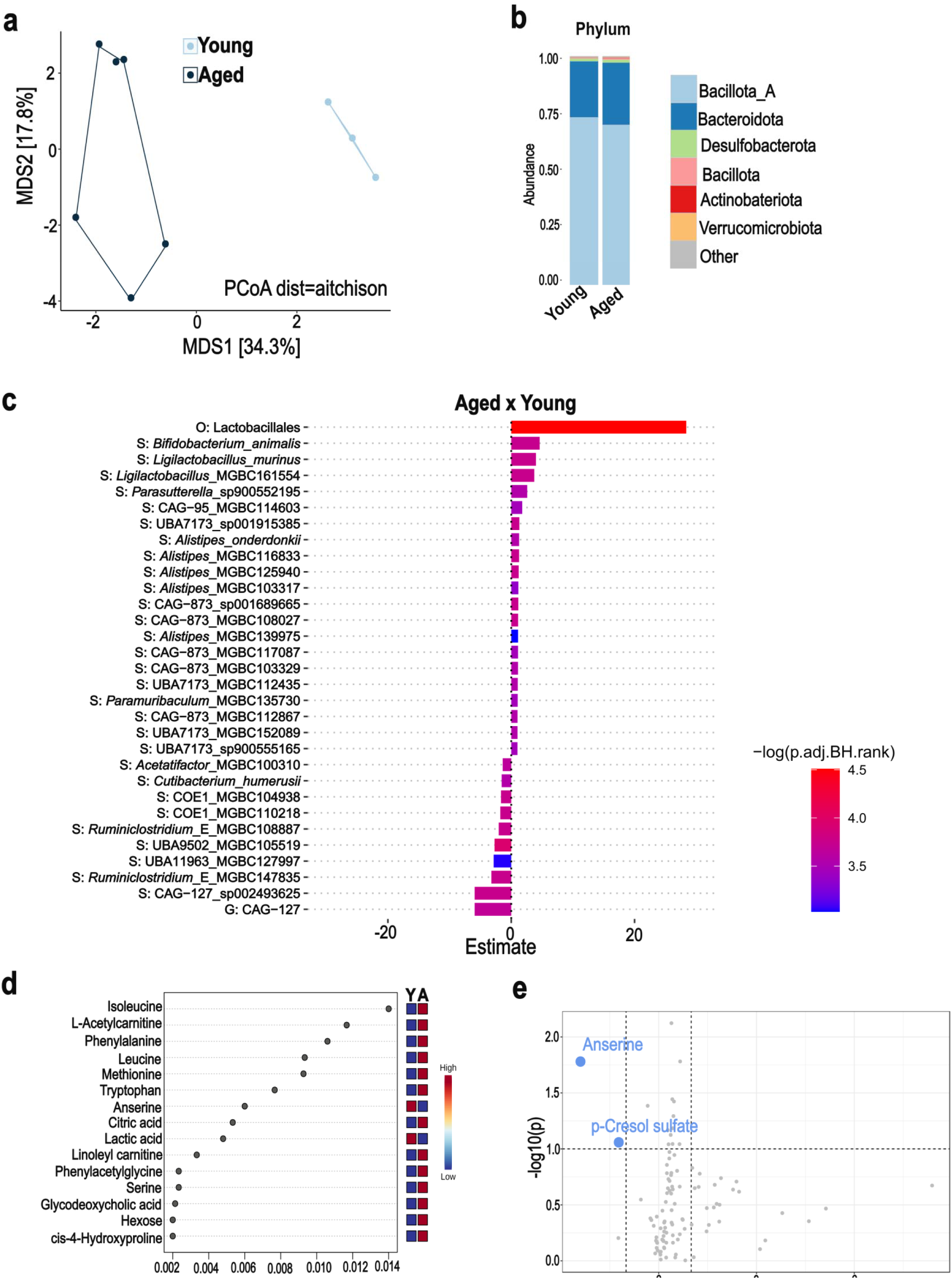
Shotgun metagenomics and plasma metabolome analysis from young and aged hamsters (steady state). **a**, PCoA plot (species) representing microbial β-diversity based on Aitchson distance metric (PERMANOVA, *P* = 0.013) **b**, Gut microbiota composition at the phylum level. **c**, Differential abundance of taxa analyzed with Maaslin2 (adjusted *P*-value < 0.05). **d**, Ranking of the 15 most important metabolites from the 104 total plasma metabolites identified by Random Forest, based on mean decrease accuracy. Colored boxes indicate relative concentrations of metabolites in each group (Young: Y, Aged: A). **e**, Volcano plot of plasma metabolite differential abundance between young and aged hamsters. The *y-*axis represents the –log10 adjusted *P* value (dashed line at α = 0.05), and the *x-*axis represents the log2 FC (dashed line at two fold change) (n=3-6/group).

Given the established link between gut microbiota and host metabolism^44,45^, we then assessed the impact of aging on plasma metabolites. Targeted quantitative metabolomic analysis was performed on plasma samples using liquid chromatography coupled with tandem mass spectrometry (LC-MS/MS). Hierarchical clustering analysis indicated differences in the metabolites concentrations between young and aged hamsters (**Supplementary Figure 1c**). Multivariate analysis with random forest^44^ identified several metabolites that discriminate between young and aged hamsters, including amino acids and their derivatives (isoleucine, phenylalanine, leucine, methionine, tryptophan, anserine, serine, and phenylacetylglycine), carnitines (l-acetylcarnitine and linoleyl carnitine), secondary bile acids (glycodeoxycholic acid) and components involved in energy metabolism (citric acid, lactic acid, and hexose) (**Figure 1d**). We also employed univariate statistical analysis to investigate differences in metabolites between aged and young hamsters. Compared to their young counterparts, aged hamsters showed a significant reduction in p-cresol sulfate, a tyrosine-derived metabolite linked to gut microbiota metabolism^46^, and anserine, a bioactive dipeptide acting through the gut microbiota^47^ (fold change threshold = 2.0, *P* < 0.05, **Figure 1e**). We conclude that at steady state (no infection), aged hamsters display notable changes in the composition and metabolic function of the gut microbiota.

### Aged hamsters have high pulmonary viral load, display a specific lung transcriptomic signature and develop post-acute sequelae

Age-related gut dysbiosis increases susceptibility to various diseases^28,48–51^ but its impact on respiratory infections remains largely unexplored. We therefore compared the outcomes of a SARS-CoV-2 infection in young and aged hamsters using a prototypic ancestral strain of the virus. Hamsters from each group were sacrificed on day 3 post-infection (D3), the peak of lung viral load; on D7, the peak of the acute phase response; and on D22, the resolving phase of infection^26,52^. Quantitative RT-PCR revealed that aged hamsters presented a significantly higher lung viral load than young hamsters on both D3 and D7 (**Figure 2a**, *left* panel). Immunofluorescence staining confirmed the high viral load in aged animals (**Figure 2a**, *right* panel and not shown). By D22, no viral transcripts were detected, indicating complete viral clearance. Compared to young hamsters, aged animals did not experience a greater body weight loss until D6 (**Figure 2b**). However, while young hamsters began to recover from D7 onwards, aged animals continued to lose body weight until D8 and did not regain their initial body weight on D22 (**Figure 2b** and **Supplementary Figure 2a**). Histopathology analysis (hematoxylin & eosin staining) showed that lung damage in terms of inflammation, hemorrhage, type 2 hyperplasia, and edema was similar in young and aged hosts on D7 (**Figure 2c**). Regarding these criteria, lung damage was strongly reduced by D22. Compared to the young group, pulmonary gene expression of C-X-C motif chemokine ligand 10 (*Cxcl10*) and interferon-stimulated gene 15 (*Isg15*) (representative of ISGs), but not chemokine ligand 2 (*Ccl2*) and interleukin-6 (*Il6*) (representative of inflammation), was higher in the aged group on D7 (**Supplementary Figure 2b**). Gene expression of the tight junction proteins zonula occludens 1 (*Zo1*) and occludin (*Ocln*) was reduced in the aged group, indicating elevated alteration of the epithelial barrier (**Supplementary Figure 2c**). To further explore the effects of aging on pulmonary gene expression, bulk RNA sequencing (RNAseq) was performed on lung tissue. This analysis revealed distinct differences in gene expression between young and aged hamsters at D7, as illustrated by Venn diagrams (**Supplementary Figure 2d**) and volcano plots (**Figure 2d** and **Supplementary Table 2**). A substantial portion of the differentially expressed genes (DEGs) was shared between young and aged hamsters (40-45%, gene ontology enrichment analysis in **Supplementary Figure 2d)**. Young and aged hamsters also had exclusive DEGs, with most being upregulated in aged hamsters (1,178/1,501) and downregulated in young hamsters (1,660/1,937). Pathways induced in infected aged hamsters included those related to protein synthesis and degradation (*e.g*., “proteasome” and “protein processing in endoplasmic reticulum”) and immune activation (“Epstein-Barr Virus infection”, “Immunity”, “Lysosome” and “NF-kappa B signaling pathway”), indicating increased protein turnover and intense immune activation (**Figure 2e**, *upper* panel). Genes exclusively downregulated in young hamsters were associated with transcription regulation and metabolism (e.g., “Propanoate metabolism”, “Valine, leucine, and isoleucine degradation”, “Lipid metabolism” pathways), as well as “Platelet activation” (**Figure 2e**, *lower* panel). Although speculative, the heightened activation of proteasome and immune pathways observed in aged hamsters may explain worse disease outcomes.

**Figure 2.**
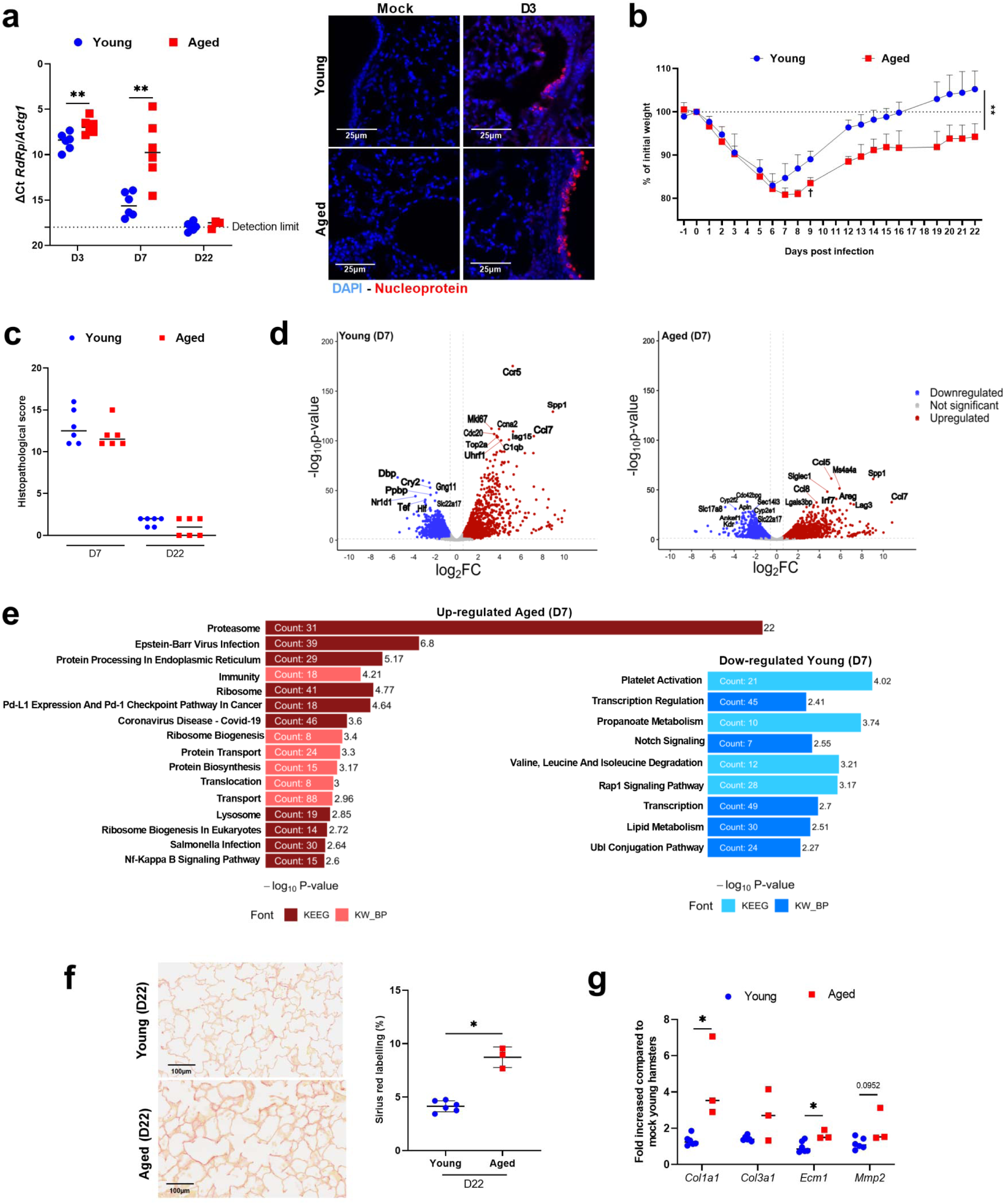
SARS-CoV-2 infection in young and aged hamsters. **a**, Lung viral load (RT-PCR) on D3, D7 and D22. *Left* panel: Quantification of viral RdRp transcript levels in lungs using RT-PCR assay. Data are expressed as the mean ΔCt. *Right* panel: Immunofluorescence staining for DAPI (blue) and viral nucleoprotein (red). Bars: 25 μm. **b**, Percentage of body weight change curves for infected animals, compared using a two tailed Mann-Whitney *U* test (area under the curve). Of note, one aged hamster lost a large amount of weight and suffered from respiratory distress at 9 dpi and so was sacrificed. **c**, Histopathological examination of lung sections (H&E staining). Total score is indicated for each group. **d**, Volcano plot of transcriptomic data generated from whole lung tissue collected from young and aged hamsters (D7). The x-axis represents the -log2 fold change for differentially expressed genes in mock vs. infected lungs, and the y-axis represents the -log10 (p.adj). Significant differentially expressed genes in infected lungs with a fold change threshold of 0.6 (vertical dashed lines) and a *P*-value < 0.05 (horizontal dashed line) are shown in red (for upregulation) or blue (for downregulation). **e**, Gene set enrichment analysis from Kyoto Encyclopedia of Genes and Genomes (KEEG) and Keywords Biological Process (KW_BP) (aged hamsters, D7). **f**, *Left* panel: Sirius Red staining images. *Right* panel: Percentage of Sirius Red labelling in lungs on D22. **g**, Expression of *Col1a1*, *Col3a1, Ecm1* and *Mmp2* gene expression in lungs from young and aged hamsters on D22 by quantitative RT-PCR. The data are expressed as the mean fold change relative to average gene expression in mock-infected young animals (6 young hamsters and 3 aged hamsters). **a-c**, **f** and **g**, n=3-6/group. **d** and **e**, n=4/group. Significant differences were determined using the Mann Whitney *U* test (**a**, **c** and **g**) (\**P* < 0.05; ** *P* < 0.01).

Clinical and experimental evidence suggests that severe respiratory viral infections can lead to persistent lung pathology, remodeling and pulmonary dysfunction^53–55^. The kinetics of lung damage after acute SARS-CoV-2 infection was then assessed through Sirius Red staining of lung sections. Aged hamsters had a significantly higher percentage of Sirius Red staining (a marker of fibrosis) on D22 compared to young animals (**Figure 2f**). Accordingly, transcript expression levels of markers associated with pulmonary fibrosis such as collagen 1 alpha 1 (*Col11*), *Col3a1*, extracellular matrix protein 1 (*Ecm1*) and matrix metalloproteinase 2 (*Mmp2*) were higher in aged hamsters than in young hamsters (**Figure 2g**). The above data pointed to an unresolved disease in aged hamsters. Altogether, young and aged hamsters respond differently to SARS-CoV-2-infection (acute and resolving phases), with a more severe and persistent disease in aged animals.

### SARS-CoV-2 infection differentially affects gut microbiota composition and function in young and aged hamsters

We then compared the gut microbiota composition and function in SARS-CoV-2-infected young and aged hamsters by analyzing cecal contents on D7 and D22. No significant changes in alpha diversity (Shannon index) were observed between the experimental groups (**Supplementary Figure 3a**). However, PCoA based on Aitchison distance revealed significant alterations in the microbiota community structure in both age groups on D7 (**Figure 3a**, *P* = 0.007 for D7 aged vs D0 aged; *P* < 0.02 for D7 young vs D0 young). On D22, the bacterial community structure shifted toward the baseline groups, but remained distinct from baseline in both aged (*P* < 0.02 for D22 aged vs D0 aged; *P* < 0.05 for D22 aged vs D7 aged) and young animals (*P* < 0.02 for D22 young vs D0 young; *P* = 0.007 for D22 young vs D7 young). On D7, aged hamsters exhibited the most pronounced shifts in microbiota composition, including a marked increase in the relative abundance of the Bacteroidota phylum (from 27% to 47%) and the Pseudomonadota phylum (from 0.05% to 3%) (**Figure 3b**, *left* panel). Within the Bacteroidota, the genera *Alistipes* and *Bacteroides* increased significantly, whereas within Pseudomonadota, the *Escherichia* genus showed a significant rise (**Figure 3b**, *right* panel and **Supplementary Figure 3b**).

**Figure 3.**
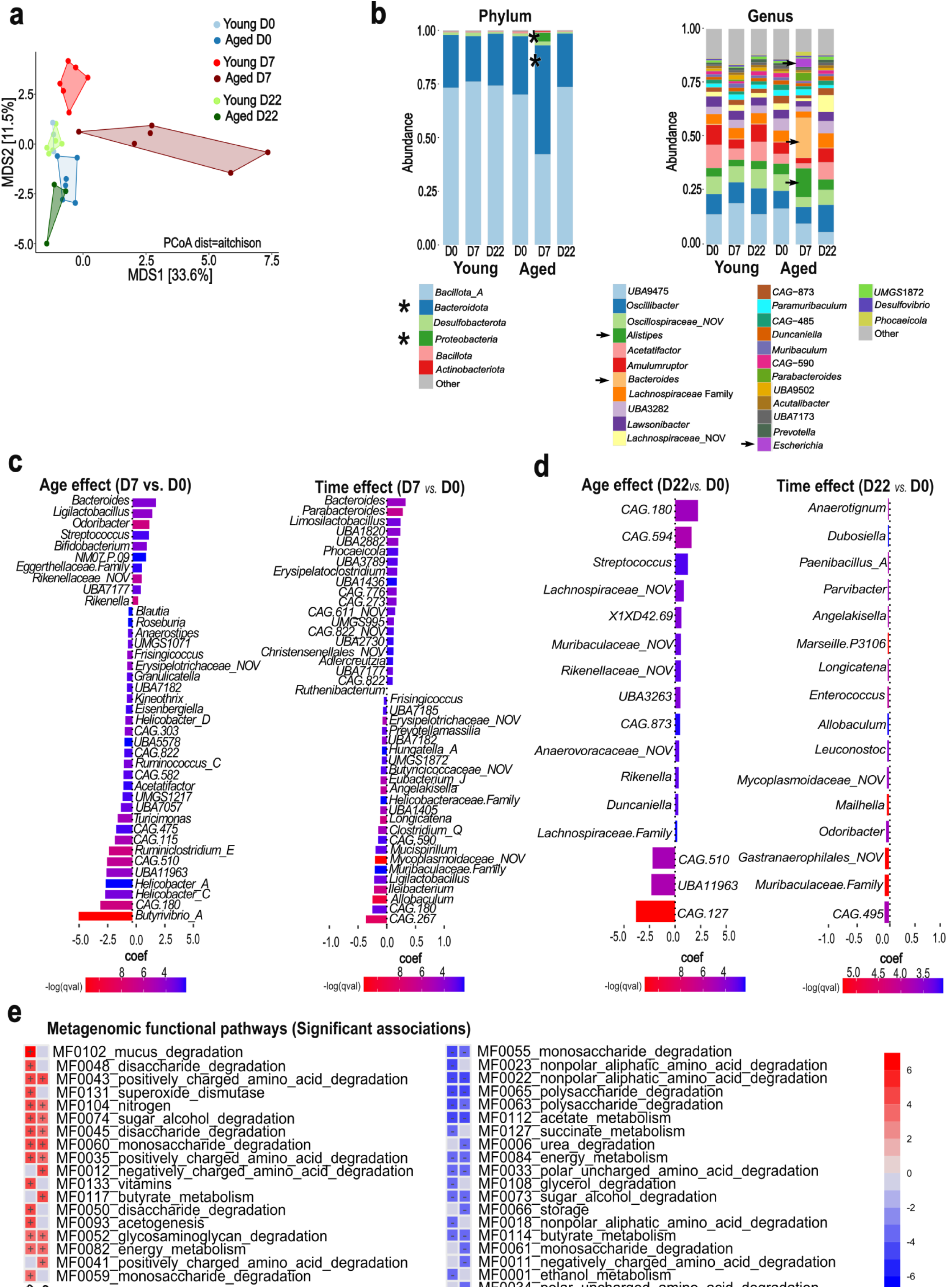
Taxonomic and functional profile of the gut microbiota across different age groups and infection states. **a**, PCoA plot (species) of β-diversity based on Aitchson distance metric, differentiating infection status (non-infected: blue, D7: red, D22: green) and age group (young: light colors, aged: dark colors). **b**, Compositional bar plots illustrating the taxonomic profile of gut microbiota at the phylum (*left* panel) and genus (*right* panel) levels. The relative abundance of each taxon is represented by the bar height. Asterisks and arrows indicate major changes between D0 and D7. **c**, MaAsLin2 multivariate differential abundance analysis for bacterial genera comparing D7 with non-infected controls, stratified by age group (young and aged). **d**, MaAsLin2 multivariate differential abundance analysis for bacterial genera comparing D22 with non-infected controls, stratified by age group (young and aged). **c-d**, *left panel*: Bar plot depicting the age effect, with positive coefficient values indicating bacterial genera that are significantly enriched in aged animals compared to young animals. *Right panel*: Bar plot depicting the time effect, with positive coefficient values indicating bacterial genera that are significantly enriched in infected (D7 or D22) animals compared to non-infected controls. **e**, MaAsLin2 multivariate differential analysis for gut microbiota functional pathways represented as a heatmap with columns for both the infection and age effects. Positive values (red) indicate enrichment in infected or aged groups.

We next employed a multivariate linear model to identify statistically significant associations between gut microbial taxa and animal metadata (age and day of infection). Between D0 and D7, aged hamsters showed significant positive associations with genera such as *Bacteroides*, *Ligilactobacillus*, *Odoribacter*, *Streptococcus,* and *Bifidobacterium* (**Figure 3c**). In contrast, the young group was enriched with genera including *Butyrivibrio_A*, *Ruminiclostridium_E*, *Ruminococcus_C*, and certain *Helicobacter* groups. For the time variable, several genera like *Bacteroides*, *Parabacteroides*, *Limosilactobacillus*, *Phocaeicola*, and *Erysipelatoclostridium,* increased on D7, whereas others such as *Allobaculum*, *Ileibacterium*, and *Ligilactobacillus* decreased. Between D0 and D22, fewer significant associations were found. Age was positively associated with the genera *CAG.180 (*f *Acutalibacteraceae*), *CAG.594* (c *Bacilli*), and *Streptococcus*, and negatively associated with CAG-127 (f *Lachnospiraceae*), UBA11963 (c *Bacilli*) and CAG-510 (f *Lachnospiraceae*) (**Figure 3d**). Time negatively associated with CAG-495 (c *Alphaproteobacteria*), *Odoribacter*, *Mailhella,* and other genera (**Figure 3d**).

To elucidate the functional implications of microbiota alterations, we conducted a multi-step metagenomic functional analysis incorporating HUMAnN^56^, GOmixer, and MaAsLin2^57^. During the acute phase of infection (D7), significant modulations in gut metabolic modules were identified, which were associated with age and infection status (**Figure 3e**). Specifically, the acute phase of infection was characterized by the upregulation of modules involved in mucus degradation, simple carbohydrate degradation, and positively charged amino acid degradation on D7. There was also an increase in modules related to nitrogen metabolism, vitamin metabolism, energy metabolism, superoxide dismutase activity, and acetogenesis (**Figure 3e)**. Interestingly, there was a downregulation of some carbohydrate degradation modules (mainly polysaccharides), non-polar amino acid degradation, and SCFA metabolism (acetate and butyrate (**Figure 3e**). These findings suggest a dynamic shift in gut microbiota function during the early stages of infection. In comparison, the resolving phase of infection (D22) showed a downregulation of carbohydrate and amino acid degradation modules (**Supplementary Figure 3c**). Conversely, there was an upregulation of modules related to ethanol and energy metabolism, indicating a shift in resource allocation or altered energy demands during recovery. The analysis of age effects also revealed contrasting patterns between the acute and resolving phases. During the acute phase, age was positively associated with charged amino acid degradation, saccharide degradation, and butyrate metabolism, and negatively associated with overall amino acid degradation, saccharide degradation (including more complex carbohydrates), urea degradation, and acetate and butyrate metabolism (**Figure 3e**). In the resolving phase (D22, **Supplementary Figure 3c**), age was negatively associated with monosaccharide degradation and SCFA metabolism, and positively associated with the degradation of more complex carbohydrates and charged amino acids. This finding suggests an age-dependent modulation of microbiota metabolic pathways during recovery.

### SARS-CoV-2 infection differentially alters plasma metabolite concentrations in young and aged hamsters

Clinical studies demonstrated the good discriminatory power for distinguishing COVID-19 disease severity based solely on a selection of circulating metabolites^23,58^. We compared the plasma metabolite levels in young and aged hamsters during SARS-CoV-2 infection. PCoA revealed that infection significantly impacted plasma metabolite concentrations on D7 in aged hamsters, with some differences persisting until D22 (**Figure 4a**). Several of these metabolites were associated with microbial metabolism^45,59^. ANOVA-Simultaneous Component Analysis (ASCA) with permutation testing revealed significant differences between experimental groups based on age (P < 0.05) and infection status (P < 0.05) (**Supplementary Figure 4a**), although no significant interaction between these variables was found. Using multivariate analysis with random forest, we identified the 15 most relevant variables for discriminating between experimental groups. This list encompassed metabolites such as cis-aconitic acid, citric acid, spermidine, and various amino acids and their derivatives, including kynurenine, creatine, glutamic acid, tryptophan, phenylalanine, proline betaine, 2-aminobutyric, serine, trans-4-hydroxyproline, and 1-methylhistidine (**Figure 4b**). Choline and the bacterial product trimethylamine (TMA) were also among the top 15 discriminating metabolites. Hierarchical clustering of the top 30 metabolites revealed two distinct clusters (**Figure 4c**). The first cluster contained metabolites with high concentrations on D7 in aged hamsters, some of which returned to baseline levels on D22 (e.g., 1-methylhistidine, kynurenine, L-carnitine, p-cresol sulfate, 2-aminobutyric acid). This cluster also included sarcosine and cis-aconitic acid, which remained elevated on D22. Notably, some metabolites within this cluster, such as cysteine, creatine, indoxyl sulfate, and spermidine, were reduced in young animals on D7. The second cluster comprised metabolites with reduced concentrations on D7 in young, aged, or both groups. Some, like citric acid and indole-3-propionic acid, returned to baseline levels on D22. However, tryptophan, phenylalanine, and glutamic acid remained low on D22, particularly in aged animals (**Figure 4c**).

**Figure 4.**
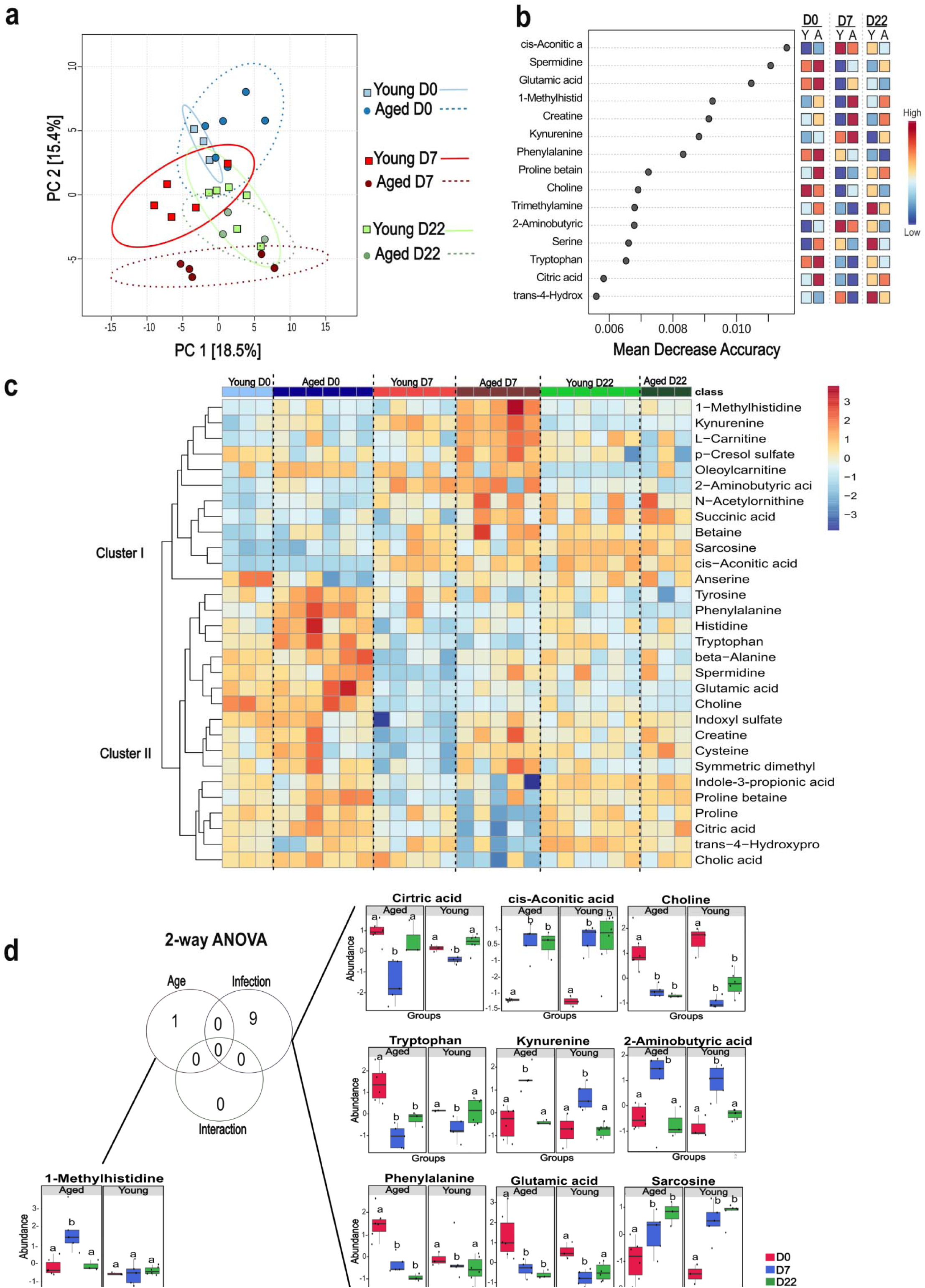
Multivariate and univariate analyses, and metabolite sets enrichment overview of plasma metabolome profiles of SARS-CoV-2-infected young and aged hamsters. Plasma metabolomic profiling of mock-infected and SARS-CoV-2-infected young and aged hamsters on D7 and D22. Shown are the multivariate analysis with PCA. [PERMANOVA] F-value: 6.3905; R-squared: 0.59223; p-value (based on 999 permutations): 0.001) (**a**) and ranking of top 15 metabolites identified by random forest, based on mean decrease accuracy (**b**). Colored boxes indicate relative concentrations of metabolites in each group, **c**, Hierarchical cluster analysis and heatmap of top 30 metabolites. Young=light blue; Aged=dark blue; infected D7 young=red and aged=dark red; infected D22 young =light green and aged=dark green. Note the elevated ratio between kynurenine (high) and tryptophan (low) at D7, a feature of the acute phase. **d,** Univariate analysis via 2-way ANOVA (threshold FDR *P* < 0.05) with Tukey’s post-test comparing infected vs. non-infected control for both age groups. Non-infected=red; infected D7=blue and infected D22=green.

We then employed a Two-Way ANOVA test within a univariate statistical framework to investigate the impact of age, infection, and their interaction on metabolite levels. Age emerged as a crucial factor contributing to the elevation of 1-methylhistidine (transient increase in aged animals) (**Figure 4d** and **Supplementary Figure 5**). Interestingly, this metabolite has been identified as a predictor of survival chances in hospitalized COVID-19 patients^58^ (**Figure 4d**). Notably, SARS-CoV-2 infection induced significant alterations in nine metabolites independently of age on D7. While citric acid, kynurenine, and 2-aminobutyric acid returned to basal levels by D22, metabolites like cis-aconitic acid, choline, and sarcosine showed sustained alterations in both young and aged groups on D22 (non-infected *vs*. D7, *P* < 0.05 and non-infected *vs*. D22, *P* < 0.05 for all age groups). Exploring metabolites associated with the prolonged effects of infection, we observed that tryptophan, phenylalanine and glutamic acid remained altered on D22 only in aged hamsters (non-infected *vs*. D22, *P* < 0.05 for the aged group) (**Figure 4d**). Using Enrichment Global Test and Topology analysis, we verified how the significant metabolites (altered after acute or long infection regardless of age) interact and connect with each other, respectively. The interaction shown in the metabolite sets enrichment overview revealed that SARS-CoV-2 infection mainly alters the metabolism of key amino acids, energetic pathways, and cell membrane components (**Supplementary Figure 4b**). The network plot further reinforced the relationship between these pathways (**Supplementary Figure 4c**). In summary, SARS-CoV-2 infection differentially modulates systemic metabolite concentrations in young and aged hamsters, revealing age-dependent metabolic responses to infection. It is noteworthy that SARS-CoV-2 infection has long-term effects in both age groups, with some changes being specific to aged hamsters.

### Microbiota taxa and metabolites associate differently with acute disease markers in young and aged hamsters

We then performed an integrative analysis combining metagenomics, metabolomics and clinical data to identify relevant associations (**Supplementary Table 3**). Using Hierarchical All-against-All (HAllA) association testing, we explored pairs of datasets, applying the Spearman correlation coefficient to measure associations for continuous data. Significant associations were visualized as networks, where each node represented a feature (metabolite, bacterial species, or infection marker) and each edge indicated a correlation between two different features. By comparing the D7 groups with non-infected controls, we generated two distinct networks for young and aged hamsters (**Figure 5**). Compared to young hamsters, aged hamsters exhibited a denser network of correlations (70 edges vs. 21 edges). In aged hamsters, amino acids, including proline, isoleucine, serine, methionine, valine, tyrosine, phenylalanine, histidine, tryptophan, and glutamic acid, showed positive correlations with body weight and/or lung barrier integrity (*Zo1*), and negative correlations with virus copy numbers, pneumonia score, *Cxcl10* and/or *Ccl2* (**Figure 5a** and **Supplementary Figure 5**). The gut bacteria species *Alistipes_sp003979135* and *D16-63_MGBC165261* (f Eggerthellaceae) were positively associated with some of these amino acids. Other metabolites, such as histamine, L-acetylcarnitine, choline, and citric acid presented similar correlation patterns. Furthermore, proline betaine and the bile acid tauromuricholic acid correlated positively with body weight, while indoleacetic acid, cholic acid, and creatine correlated negatively with the pneumonia score. Indoleacetic acid also correlated negatively with virus copy numbers (**Figure 5a**). Associations between bacterial taxa and metabolites were also observed in the aged group on D7, although we noticed a lack of direct correlation between taxa and disease markers. *Bacteroides tethaiotaomicron* and proline betaine showed a negative correlation, while *UBA1394* (f Ruminococcaceae) was positively correlated with L-acetylcarnitine, and three other UBA11940 (f Borkfalkiaceae) species were positively correlated with indoleacetic acid (**Figure 5a**). In young hamsters, a distinct profile emerged, with virus copy number correlating negatively with carnosine, beta-alanine, and *COE1_sp002358575* (*Lachnospiraceae* family) (**Figure 5b**). Another species of the *Lachnospiraceae* family, *COE1_MGBC117419*, and carnosine, correlated negatively with pneumonia score. Furthermore, cis-aconitic acid correlated negatively with body weight and *Zo1*, the later positively associated with *Eubacterium, Lawsonibacter* and *Oscillospiraceae*. Body weight also presented negative correlation with *Turicimonas muris* and positive correlation with *Lawsonibacter_MGBC000538* and *UBA7182_MGBC119923 (*f Lachnospiraceae (**Figure 5b**). Overall, our findings reveal distinct metabolic and microbial associations during SARS-CoV-2 infection between young and aged hamsters on D7.

**Figure 5.**
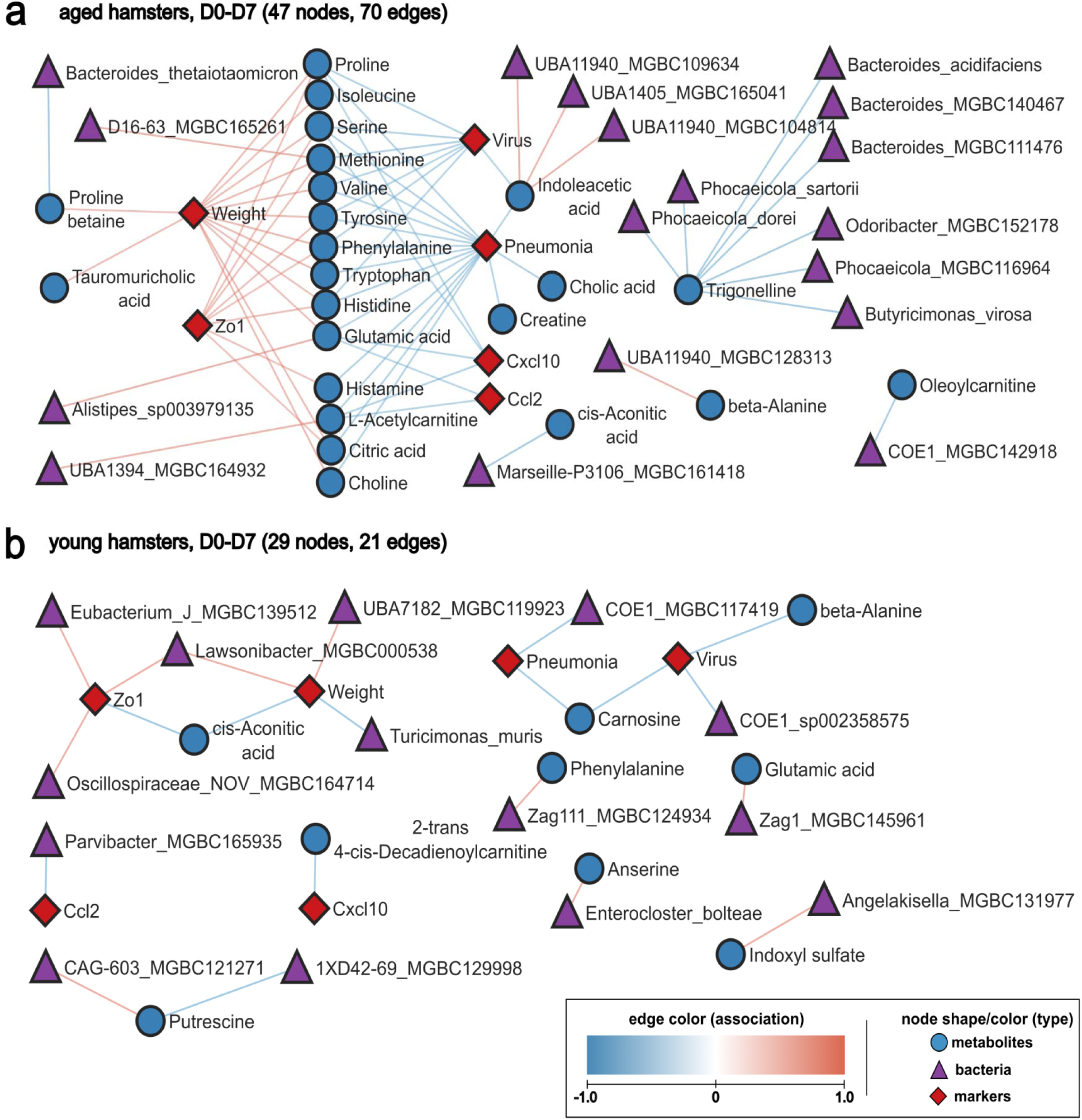
Correlation between systemic metabolites, taxa and early infection-related variables (D0 *vs* D7). **a** and **b**, Continuous data from hamsters infected with SARS-CoV-2 (D7 and non-infected controls) were used to calculate Spearman correlation coefficient, with adjusted *P*-value ≤ 0.05. Each node represents a biological entity (metabolites, gene, bacterial species, or infection markers). Edges represent statistically significant associations between the connected nodes of different types, with red edges for positive and blue edges for negative correlation. **a,** Network for aged hamsters. The correlation network centered around amino acids and some of their derivative molecules with positive association with body weight and negative association with viral load and inflammation. **b,** Network for young hamsters.

### Amino acids and specific metabolites emerged as key indicators of disease progression in aged animals

Aging favors non-resolving damage following SARS-CoV-2 infection, including delayed body weight recovery (**Figure 2b** and **Supplementary Figure 2a**) and development of lung fibrosis (**Figure 2f** and **g**). We therefore analyzed potential correlations between these disease markers, metabolites and microbiota changes on D22. For that purpose, we compared the D22 group to non-infected controls and generated distinct networks for young and aged hamsters (**Figure 6**). In aged hamsters, body weight correlated positively with several amino acids (tyrosine, tryptophan, phenylalanine, glutamic acid and methionine), as well as with choline, citric acid and tauromuricholic acid (**Figure 6a** and **Supplementary Figure 6**). Conversely, fibrosis correlated negatively with methionine, histidine, homoarginine, creatine and indoleacetic acid. Other metabolites like indoxyl sulfate, 1-methylhistidine, cholic acid, valine, lysine, beta-alanine, indoleacetic acid and glutamic acid correlated negatively with the fibrosis marker genes *Mmp2*, *Ecm1* and/or *Col1a1* (**Figure 6a**). It is noteworthy that on D22, no bacterial taxa associated with body weight and fibrosis in aged animals. In the young group, some bacterial species (from the *Paramuribaculum* and CAG485 genera) and metabolites (anserine) correlated negatively with fibrosis, suggesting a possible role of these components in the recovery phase. Apart from that, limited correlation between bacterial species and/or metabolites and disease markers were observed in young animals on D22. Overall, some amino acids and specific metabolites emerged as key indicators of disease progression in aged animals, with strong associations with body weight and fibrosis.

**Figure 6.**
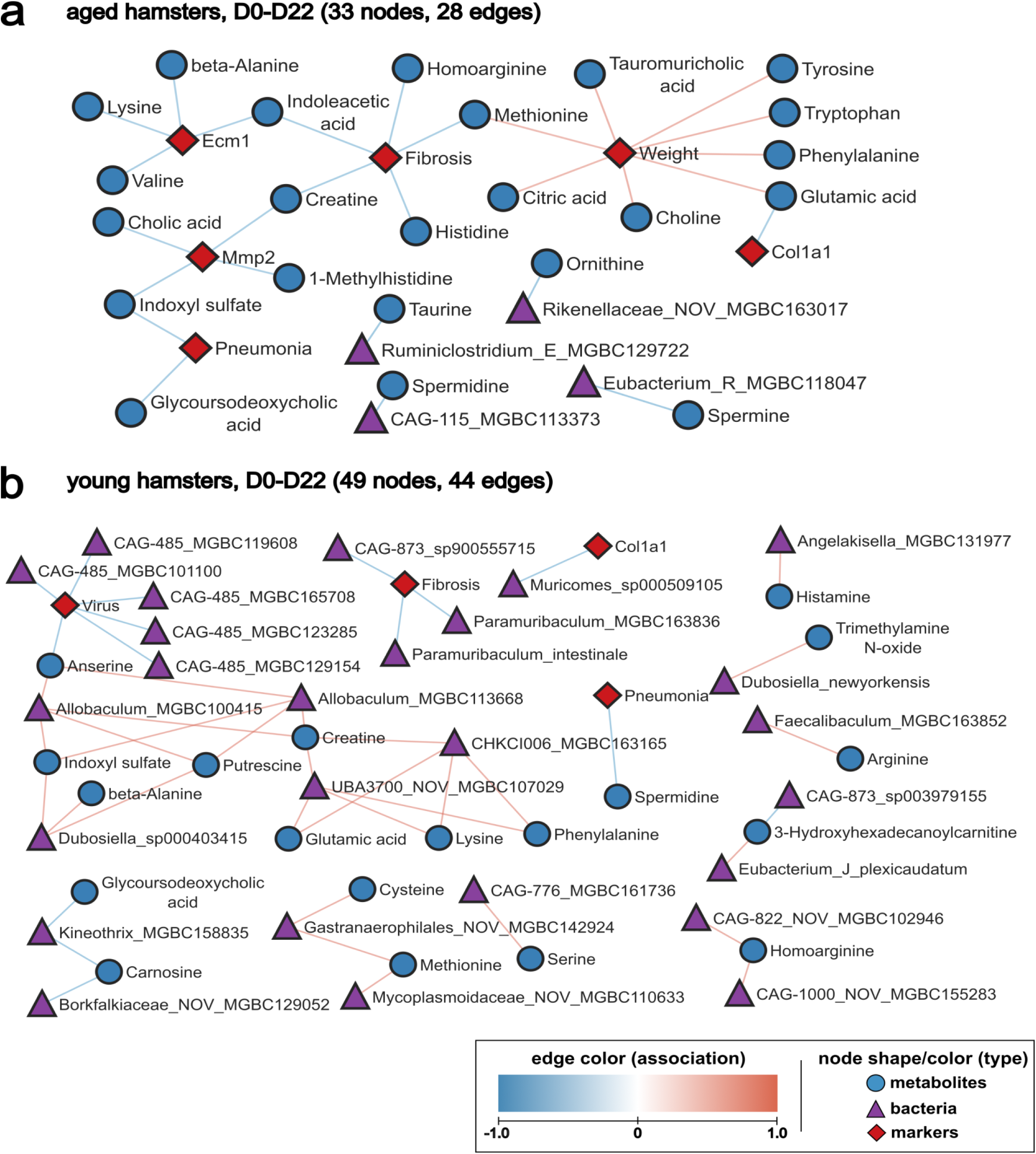
Correlation between systemic metabolites, taxa and late infection-related variables (D0 *vs* D22). **a,** Network for aged hamsters. **b,** Network for young hamsters. **a** and **b**, Continuous data from hamsters infected with SARS-CoV-2 (D22 and non-infected controls) were used to calculate Spearman correlation coefficient, with adjusted *P*-value ≤ 0.05. Each node represents a biological entity (metabolites, gene, bacterial species, or infection markers). Edges represent statistically significant associations between the connected nodes of different types, with red edges for positive and blue edges for negative correlation.

## Discussion

Clinical studies have evidenced changes in gut microbiota composition during SARS-CoV-2 infection^9,11–14,20,22,23^, this alteration can persist for many months^10,16–19,21,24^. However, metagenomic characterization for advanced age individuals remains elusive. Since clinical studies typically involved hospitalized patients, it is challenging to distinguish between the direct effects of the viral infection and the impacts of hospitalization and medication on the metagenomics results. Therefore, preclinical models are instrumental to analyze the potential impact of advanced age on virus-induced dysbiosis and on subsequent disease outcomes. In our study, we used high-throughput shotgun metagenomics and metabolomics analyses to identify potential bacterial and metabolite markers of acute and long-term COVID-19 in aged hamsters, revealing the impact of aging on gut microbiota composition and disease outcomes.

In the absence of infection, we found altered microbiota composition in 22-month-old hamsters (∼80 years old for humans), as indicated by beta diversity analysis. Aged hamsters showed enrichment in the order *Lactobacillales* and species including *Bifidobacterium animalis* and *Ligilactobacillus murinus*, and *Alistipes* genus. *Lactobacillales* and *Bifidobacterium* species are abundant in healthy elderly individuals^60,61^. In mice, *B. animalis* is positively associated with longevity^62^ and/or good health^63,64^. The *Alistipes* genus has also been reported to increase in the gut microbiota of aged individuals, despite some controversial observations^28,65^. These findings suggest that our aged non-infected hamsters may represent a model of healthy senile gut microbiota. Regarding systemic metabolic output, anserine and p-cresol sulfate levels were reduced in the plasma of aged hamsters compared to young hamsters. Anserine, a natural carnosine derivative, possesses anti-inflammatory and antioxidant properties that are dependent on gut microbiota^47^. The decline in anserine levels with age may be due to multiple factors, including changes in the digestion and absorption of dietary nutrients^66^, as seen in aged mice ^67^. Anserine supplementation has been linked to improvements in cognitive and inflammatory processes during aging^68,69^. p-cresol sulfate is the fermentation product of p-cresol, a gut microbiota-derived product of L-tyrosine^46^. It also has anti-inflammatory properties, including in the lungs^70^. To the best of our knowledge, its concentration in aged plasma (in rodents and others) has not yet been described. Based on these findings, and in line with observations in other species^28^, aged hamsters exhibit alterations in both the composition and functionality of the gut microbiota under steady-state conditions. These changes may play a role in shaping the host response to infection. Indeed, compared to young animals, aged hamsters showed higher viral loads and experienced unresolved disease (more intense and prolonged weight loss and greater pulmonary fibrosis) following SARS-CoV-2 infection. Whether pre-existing gut dysbiosis distally affects lung defence, body weight recovery and/or lung fibrosis in aged individuals remains to be demonstrated. Further experimental exploration, including fecal transfer experiments, is required to investigate this issue.

The influence of the age on virus-induced gut dysbiosis was then assessed. Our data show that SARS-CoV-2 infection elicited more pronounced alterations in the gut microbiota of aged hamsters compared to young hamsters. In aged hamsters, the genera *Bacteroides*, *Ligilactobacillus*, *Odoribacter*, *Streptococcus*, and *Bifidobacterium* increased while in the young group, *Butyrivibrio_A*, *Ruminiclostridium_E*, *Ruminococcus_C*, and certain *Helicobacter* groups were more abundant. Taxonomic changes were relatively transient, although some genera showed persistent alterations in the aged group. These include the detrimental genera *Streptococcus* (increased) and members of the beneficial, SCFA producer family *Lachnospiraceae* (CAG-127 and CAG-510) (decreased). More research is also required to better understand the role of these species in different phases of COVID-19 such as the acute inflammatory phase and the subsequent periods of resolution and tissue repair. To address this gap, fecal transfer experiments would be necessary. It is noteworthy that this approach is instrumental to study the consequences of virus-induced dysbiosis on the early and later outcomes of respiratory viral infections^71,72^. Functional analysis of metagenomics data revealed that several microbial metabolic pathways were altered during infection, many of which were also affected by aging. Notably, there was an increase in pathways related to energy metabolism, degradation of monosaccharides and disaccharides, and degradation of positively charged amino acids. At the same time, pathways involved in polysaccharide degradation were reduced. Both aged and young hamsters showed a reduction in SCFA (acetate and butyrate) production pathways after infection. This is in line with the reduced frequencies of SCFA producers such as *Butyrivibrio_A*, *Ruminiclostridium_E*, and *Ruminococcus_C* in infected aged animals (the current study). Reduced SCFA production during acute respiratory infection has been reported in preclinical and clinical studies^16,26,37,71,73–75^. These gut microbiota functional alterations induced by infection, along with other age-associated changes, may contribute to the worsened response to SARS-CoV-2 infection in aged individuals. Aging plays a significant role in the long-term impact of infection. At D22, aged hamsters showed a negative association with butyrate and acetate metabolism pathways. Reduction of SCFA metabolism is a hallmark of aged individuals^28^ and a marker of long COVID-19 in humans^75^. Moreover, and in line for our metabolic analysis, age was positively associated with the degradation of amino acids during the acute and recovery phases of infection. Together, our metagenomics analysis demonstrates that SARS-CoV-2 infection triggers a pronounced perturbation of the gut microbiota’s composition and function in aged individuals. Of note, this finding was recently confirmed using a metaproteomics approach (Creskey and *al*., submitted).

SARS-CoV-2 infection significantly impacts on metabolite levels in both young and aged hamsters, but the types of metabolites and their clustering patterns, associated with microbiota composition and disease markers, differed between the age groups. For example, aged hamsters exhibited a transient elevation in 1-methylhistidine, a product of histidine methylation associated with muscle protein turnover and metabolism^76^, body weight loss, and sarcopenia^77^. As metabolic disorders and sarcopenia significantly affected hospital outcomes and increased the risk of physical decline post-COVID-19 recovery^78–80^, systemic 1-methylhistidine in patients might serve as a valuable prognostic factor^58^. By integrating our various omics data, we identified associations between the gut microbiota, metabolome, and infection markers. Several amino acids (phenylalanine, tryptophan, glutamic acid) and indoleacetic acid emerged as potential protective factors during both the acute and resolving phases of SARS-CoV-2 infection in aged hamsters. For instance, the sustained drop of tryptophan, phenylalanine, and glutamic acid correlated with the lack of body weight recovery in aged animals on D22. In severe COVID-19 cases, patients in intensive care units show increased degradation of these amino acids due to heightened gluconeogenesis and lipid breakdown^81,82^, which is associated with COVID-19 outcomes^19^. More studies are needed to investigate the pathways involved in these changes and to identify potential treatment targets to reduce the impact of infection on amino acid metabolism and protein turnover. Our data suggest the importance of prolonged monitoring of elderly individuals, especially regarding nutritional support of amino acids. The microbiota-derived tryptophan metabolite indoleacetic acid also negatively associated with key disease markers at D7 and D22 in aged animals. This metabolite is well described for its anti-inflammatory and tissue protective role, including in the lungs^83–85^. It is noteworthy that reduced indole-3-propionic acid, another microbial tryptophan metabolite, also correlated with disease outcome during viral pneumonia^19,37,86^. In young hamsters, carnosine may act as a potential protective factor, while cis-aconitic acid appears as a potential acute disease marker on D7. In the aged group, bacterial taxa did not directly correlate with disease markers on D7 and D22. However, based on indirect associations, some gut microbiota components such as *Alistipes_sp003979135* (glutamic acid), *D16-63_MGBC165261* (f *Eggerthellaceae*) (methionine) and UBA11940/UBA1405 (f *Borkfalkiaceae*) (indolacetic acid) may be beneficial during the acute phase. In young hamsters, different species including members of *Eubacterium*, *Lawsonibacter*, *Oscillospiraceae*, COE1 (f *Lachnospiraceae*), and UBA7182 (f *Lachnospiraceae*) were associated with disease severity and may provide protection on the acute phase of infection in young hamsters. Other species such as *Paramuribaculum* and CAGs may also be protective, but on the resolving phase of the infection (D22).

Our study is novel in that we experimentally assessed gut microbiota changes and their associations with systemic metabolites, disease severity markers across two different ages and at different time points post-infection (acute and recovery phases). The current study provides several links between aging, gut microbiota components, metabolites, and the response to infection and supports the development of age-adapted, personalized microbiome-targeting strategies (nutritional, pre/pro/postbiotics) to manage disease outcomes. However, the study also had some limitations. First, the hamster model does not replicate all the features of severe COVID-19; the hamster rapidly resolves the COVID-19-like disease, although post-acute sequelae persist^52,87^. Moreover, compared to severe COVID-19 patients, the hamster model is less relevant when referred to gut disorders. Another limitation that must be acknowledged pertains to our metabolomics analysis, which, due to the large volume of plasma required, did not include SCFAs. Given the key role of SCFAs in health and disease and the impact of respiratory viral infections on SCFA levels^26,39^, SCFAs might correlate with disease markers in aged hamsters, as suggested in humans^16,75^. Thirdly, while our study provides insights into potential protective metabolites and bacterial taxa, correlation does not imply causation, warranting cautious interpretation. Functional studies are therefore requested to experimentally test the potential effects of candidate metabolites, such as certain amino acids, indoleacetic acid and 1-methylhistidine, in the COVID-19-like severity. Despite these limitations, our study provides the first description of gut microbiota changes during experimental severe COVID-19 in aged hamsters. Identifying factors contributing to the severity of COVID-19, as well as other acute respiratory viral infections such as influenza, in elderly individuals is a first step towards developing more personalized therapies for this condition in order to reduce the risk of death and lower long-term sequelae.

## Material and Methods

### Animals and ethics statement

Golden Syrian hamsters were purchased from Janvier Laboratory (Le Genest-Saint-Isle, France). Hamsters aged 2 months (young group) or 22 months (aged group) were provided with standard rodent chow (SAFE® A04, Augy, France) and water *ad libitum* throughout experiments. All animal protocols were reviewed and approved by the local ethics committee “Comité d’Ethique en Expérimentation Animale” (CEEA) Nord/Pas-de-Calais 75. The study was authorized by the “Education, Research and Innovation Ministry” under registration number APAFIS#25041-2020040917227851 v3.

### Infection with SARS-CoV-2

Experiments involving live SARS-CoV-2 were conducted in the biosafety level 3 laboratory (BSL3) within the Institut Pasteur de Lille, adhering to national and institutional regulations and ethical guidelines (Institut Pasteur de Lille/B59-350009). After a 7-day acclimation in isolators within the BSL3 facility, hamsters were anesthetized with intraperitoneal injections of ketamine (100 mg/kg), atropine (0.75 mg/kg), and valium (2.5 mg/kg). They were then intranasally infected with 100 μl of DMEM, containing either 2 × 10^4^ TCID50 of SARS-CoV-2 (BetaCoV/France/IDF/0372/2020) for infected animals or DMEM alone for non-infected control animals. Body weight was monitored before and during infection. For tissue collection, animals were euthanized with an intraperitoneal injection of euthasol (140 mg/kg). Blood, lungs, and feces were collected from non-infected (day 0 (D0)) and SARS-CoV-2-infected hamsters on D3, D7, and D22 (3 to 6 animals per group).

### Quantification of viral load and determination of gene expression by quantitative RT-PCR

Lung tissues were homogenized in 1 ml of RA1 buffer containing 20 mM of Tris(2-carboxyethyl)phosphine hydrochloride, and total RNAs were extracted following the manufacturer’s recommendations (NucleoSpin RNA kit, Macherey-Nagel). For the determination of gene expression, RNA was reverse-transcribed using a High-Capacity cDNA Archive Kit (Life Technologies, Carlsbad, CA, USA). The resulting cDNA was amplified using SYBR Green-based real-time PCR and the QuantStudio™ 5 K Flex Real-Time PCR System (Applied Biosystems). Viral RNA was quantified using qRT-PCR with SYBR green PCR master mix (Life Technologies, Carlsbad, CA) and primers for RNA-dependent RNA polymerase (*RdRp*) fragment amplification^51^ (**Supplementary Table 4**). Individual mRNAs were quantified relative to the expression of genes coding for RdRp and gamma actin (Actg1). The viral load was expressed as the amount of viral RNA relative to *Actg1* expression level (ΔCt). For host gene expression, relative mRNA levels were determined according to the 2^-ΔΔ^Ct (cycle thresholds) method by comparing (i) the PCR Ct for the gene of interest and the housekeeping gene (ΔCt) and (ii) the ΔCt values for the treated and control groups (ΔΔCt). Data were normalized against the expression of the *Gapdh g*ene and expressed as fold-change over the mean gene expression level in mock-treated mice. Specific primers are shown in **Supplementary Table 4**). To detect SARS-CoV-2 in lung sections, lung tissue sections (7 µm thick) were dried for 48 h at 42 °C. Slides were rehydrated with toluene (AnalaR NORMAPUR ACS, VWR) and decreasing concentrations of ethanol in water. The rehydrated tissue sections were first treated with antigen unmasking solution (sodium citrate buffer pH 6). Then, sections were rinsed and blocked for 3 h at room temperature in blocking solution (PBS containing 5% BSA and 0.3% Triton X-100). To detect SARS-CoV-2, lung sections were incubated overnight at 4 °C with SARS-CoV nucleocapsid antibody (NB100-56576, 1:500, Bio-Techne) diluted in blocking solution. Sections were then washed and incubated at room temperature for 1 hour with Alexa Fluor-conjugated secondary antibodies (A-11037, 1:500) in the blocking solution. The coverslips were then mounted on slides using fluorescence mounting medium conjugated with DAPI (DAPI Fluoromount-G® #0100–20, SouthernBiotech, Birmingham, Al). Mounted slides were stored in the dark and at 4°C until image acquisition.

### Histopathological assessments

Lung tissues were fixed in 4% PBS-buffered formaldehyde for 7 days, rinsed in PBS, transferred into a 70% ethanol solution, and processed into paraffin-embedded tissue blocks. The histological processing and analysis was subcontracted to Sciempath Labo (Larçay, France). Histopathologic scores were given by a board-certified pathologist. Tissue sections (3 µm-thick) were stained with hematoxylin and eosin (H&E) reagent. Whole-mount tissues were scanned with a Nanozoomer (Hamamatsu photonics, Hamamatsu, Japan), and morphological changes were assessed by using a semi-quantitative dual histopathology score adapted from^88,89^. Different parameters were included to determine the histological scores. To evaluate pulmonary fibrosis, the Sirius Red-stained areas on scanned sections were measured with a computer-assisted, automated, whole-section histomorphometric image analysis technique (Visiopharm, Hørsholm, Denmark). Virtual whole sections were observed at a magnification of 20x (corresponding to 0.46 μm/pixel). An algorithm for Sirius Red morphometric measurement on lung-stained sections was generated with the Bayesian linear segmentation tool in the Visiopharm software package (Hørsholm, Denmark) and then refined by training on a subset of lung sections. The Sirius-Red-positive area (in mm^2^) was measured without major histology section artefacts (large vascular and peribronchiolar structures, and the alveolar lumen) and expressed as a percentage of the total area of interest. The accuracy of the automated morphometric evaluation was checked on each individual image.

### RNA sequencing analysis and statistical analysis

RNA quality was evaluated by spectrophotometry (Nanodrop, Thermo Fisher), RIN (Bioanalyzer 1200, Agilent Technologies), and quantified by fluorimetry (Qubit, Thermo Fisher). Libraries were prepared from 200 ng of total RNA using the QIAseq stranded mRNA library kit (Qiagen), according to the manufacturer’s protocol. Libraries were controlled using the Bioanalyzer 1200 (Agilent Technologies) and quantified by quantitative PCR (KAPA Library Quantification Kit for Illumina platforms, KapaBiosystems). Libraries were normalized and pooled equimolarly before sequencing on a NovaSeq sequencer (Illumina) in 2 × 150 bp mode. Raw sequencing datasets were analyzed using the nf-core/rnaseq v3.6 pipeline^90^. Raw FastQ files were quality and adapter trimmed with Trim Galore v0.6.7. (https://www.bioinformatics.babraham.ac.uk/projects/trim_galore/). Ribosomal RNA reads were filtered with SortMeRNA v 4.3.4. Cleaned reads were aligned with STAR v2.6.1d^90^ against the MesAur1.0 genome (GCA_000349665.1) and gene expression quantified using Salmon v1.5.2^91^ in mapping-based mode. Secondary analysis was performed with SARTools v1.7. 4^92^. Samples from 2 to 4 independent hamsters per group were analyzed. Counts were normalized using the trimmed mean of M values method, and differential expression analysis was done with EdgeR v3.34.1^92^. P-values were adjusted by the Benjamani-Hochberg procedure, with features filtered based on an adjusted P-value threshold of 0.05. Volcano plots were created using the tidiverse and ggrepel packages to depict the log of adjusted P-values against the log ratio of differential expression. Differentially expressed genes or gene lists of interest were input into the Database for Annotation, Visualization, and Integrated Discovery (DAVID) for enrichment analysis related to Biological Processes and KEGG pathways.

### Genomic DNA extraction and shotgun sequencing

Cecal samples were collected from young and aged non-infected (sacrificed 7 days after DMEM inoculation, termed D0) and SARS-CoV-2-infected (D7 and D22) hamsters. Cecum homogenates were stored at -80°C until analysis. Microbial DNA was extracted from approximately 150 mg of cecal samples (ZymoBIOMICS DNA Microprep Kit, Ozyme). Genomic DNA was purifiedwith Agencourt AMPure XP magnetic beads (Beckman Coulter, Brea, CA) and quantified. Libraries were prepared from 17.5 ng of genomic DNA (DNA fragment and library prep kit, sparQ, QuantaBio, Beverly, MA, USA). Libraries pooled equimolarly at 2 nM concentration. Shotgun sequencing was performed using a 150-bp paired-end sequencing protocol on an NovaSeq 6000 platform (Illumina).

### Metagenomic analysis

Raw metagenomics sequencing data were quality controlled using FastQC and MultiQC^93^. KneadData^94^ was employed to remove human and host reads (*Homo sapiens*, GCF_000001405.40 and *Mesocricetus auratus*, GCF_017639785.1), adapters, and overrepresented sequences. Taxonomic classification was performed with Kraken 2, a system based on exact k-mer matches that inform a classification algorithm^95,96^ using the database provided in the integrated Mouse Gut Metagenomic Catalog (iMGMC)^97^ as reference, with the parameter --minimum-hit-groups set to 2 and the others kept as default. Then, abundances were estimated at the species level with Bracken (Bayesian Reestimation of Abundance with KrakEN), setting the read length parameter to 150. Taxonomy results were analyzed and visualized with R libraries phyloseq^98^ and microViz (https://doi.org/10.21105/joss.03201). Taxonomy barplots were generated with the comp_barplot() function in microViz. Alpha-diversity was calculated with the ps_calc_diversity() function, using the Shannon index as measure. Beta-diversity Aitchison metric was calculated with dist_calc(), Principal Coordinate Analysis visualization was obtained with ord_calc(), and Permanova statistics was calculated with multilevel pairwise comparison using pairwiseAdonis. Statistical analysis of differential abundant taxa was conducted with Microbiome Multivariable Association with Linear Models (Maaslin2)^57^, with the parameters transform = "NONE", analysis_method = "CPLM", min_abundance = 0.01, max_significance = 0.05, and the others set as default. The HMP Unified Metabolic Analysis Network (HUMAnN) 3.0 software^56^ was employed to evaluate the abundance and functional potential of the microbial communities. HUMAnN utilized the taxonomic profile generated by Kraken 2 and Bracken, and employed the ChocoPhlAn and UniRef databases for nucleotide and protein sequences mapping, respectively. KOs (KEGG Orthology) abundances obtained from HUMAnN 3.0 were uploaded to the gut-specific metabolic pathway analysis tool, GOmixer v1.7.5.0 (Raes Lab, Ghent, Belgium), for quantification of gut metabolic pathway modules. GOmixer employed a scaling method of multiplication by a scaling factor, utilized the median as the abundance estimator, and performed an automatic guess for the minimum module coverage cutoff. Statistical comparison of the identified gut metabolic pathway modules between groups was performed using MaAsLin2. The analysis employed a log transformation (transform = "LOG"), linear model (analysis_method = "LM"), and a significance threshold of α = 0.05. Default settings were used for all other parameters.

### Metabolomic analysis

A targeted quantitative approach was implemented to analyze the hamster plasma samples. This method was based on the MxPⓇ Quant 500 kit (Biocrates Life Sciences AG, Innsbruck, Austria) using Flow Injection Analysis (FIA) and LC-MS/MS^37^. This kit quantifies 630 metabolites from three main subclasses: lipids (523), hexose (1), and small molecules (106). This technique uses isotope-labeled internal standards and provides quantitative results based on calibration curves and rigorous quality control analysis (QCs). Briefly, 10 μL of plasma samples were loaded onto a a filter containing internal standards for normalization. The filters were dried using a nitrogen stream and pressure manifold, and then incubated with phenyl isocyanate derivatization reagent for 60 minutes. After nitrogen drying, analytes were extracted with 5 mmol/L ammonium acetate in methanol and diluted for UPLC-MS/MS analysis (supplied by Biocrates). The analysis was performed on a QTRAP 5500 System (Sciex, Framingham, MA, USA) with an FIA method or coupled to a UFLC-20XR (Shimadzu, Kyoto, Japan) using a column provided in the kit. Multiple reaction monitoring was used for the quantification. MetIDQ software (Biocrates) was used to calculate the concentrations of individual metabolites. The experiments were validated using calibration curves and quality-control protocols. For each metabolite, the peaks were quantified using the area under the curve. Metabolites containing more than 10% of missing values per group (< LLOQ, < LOD) were discarded. Missing values (one value maximum out of eight) were replaced with the median of the respective groups. The data filter excluded metabolites with more than 60% of missing values. For two metabolites, ornithine, and taurine, the missing values for some samples (aged mock number 4 for ornithine and samples from young mock number 2, aged mock number 5, and aged D7 number 1 for taurine) were replaced by the mean values of their respective group. Regarding data processing, the concentration (µM) raw data was normalized by sum, log-transformed and autoscaled (mean-centered and divided by the standard deviation) and statistical analysis were performed by MetaboAnalyst 5.0^99^. Several analytical approaches were used to compare the experimental groups: Principal Component Analysis (PCA) was conducted using Euclidean distance, followed by Hierarchical Clustering with average linkage using centroid-based similarity. The resulting clusters were visualized in a heatmap. Random Forest Classification was employed to rank features by their contribution to classification accuracy, as measured by Mean Decrease Accuracy. For examining individual differences, Univariate analysis, showcasing fold changes with volcano plots specifically for non-infected young and aged hamsters was utilized. Two-way ANOVA, followed by Tukey’s HSD post hoc test for further validation was used to assess the influence of age and infection on the plasma metabolome.

### Correlations network analysis

For correlation analysis, metabolomics data was pre-processed by filtering the 53 modulated metabolites identified by random forest (Supplementary Table 3). This list identifies the features that most significantly influence the model’s performance, thus facilitating the prioritization of metabolites for subsequent analyses. For metagenomics data, species-level relative abundance values were used for all groups. For early infection (D7 vs. D0) correlation analysis, we selected infection parameters including body weight, pneumonia score, viral load, and lung gene expression as determined by quantitative RT-PCR. For the sake of simplicity, representative genes were chosen including *Cxcl10* (ISG), *Ccl2* (inflammation), and *Zo1* (barrier integrity). Late infection analysis (D22 vs. D0) additionally incorporated fibrosis score and expression of representative genes associated with fibrosis including *Col1a1*, *Col3a1*, *Mmp2*, and *Ecm1*. Hierarchical All-against-All association (HAllA)^100^ was employed with the following parameters: Spearman’s rank correlation coefficient as the similarity metric, Benjamini-Hochberg method for controlling the false discovery rate (FDR) at α = 0.05, and default expected false negative rate (FNR) of 0.2 for detecting densely associated blocks. The significant associations were then integrated into a single table and imported into Cytoscape^101^ for network visualization and analysis.

### Statistical analyses

Except for the transcriptomics, metagenomics and metabolomics analyses, all statistical analyses were performed using GraphPad Prism v9.2.0 software. Due to the low number of replicates, the normality of the data could not be tested. A two tailed Mann-Whitney *U* test was used to compare two groups. Comparisons of more than two groups with each other were analyzed with the one-way ANOVA Kruskal-Wallis test (non-parametric), followed by Dunn’s post-hoc test. Except for the transcriptomic analysis, data are expressed as the mean ± Standard Error of the Mean (MEM) or Standard Deviation (SD).

## Abbreviations

SARS-CoV-2: severe acute respiratory syndrome coronavirus 2
COVID-19: coronavirus disease 2019
SCFAs: short-chain fatty acids
dpi: day post-infection
PCoA: principal coordinate analysis
RT-PCR: reverse transcription polymerase chain reaction
MaAsLin2: Microbiome Multivariable Association with Linear Models 2

## Author contributions

FT and MARV conceived and supervised the study. FT designed the experiment. VS and FSA performed animal experiments. PBR, AD, JH, JFG and IW supervised the metabolomics analysis, and VRR, NB, HS, and MRV, supervised the gut microbiota composition analysis. DH performed the RNAseq analysis. PBR, VRR, NB, MARV and FT analyzed the data. LD and CR performed the IHC. PBR, VRR, LD, MRV and FT designed the figures. PBR, VRR, MARV and FT: drafted the manuscript. All authors revised the manuscript and provided critical comments. FT obtained funding.

## Additional informations

### Fundings

This work was supported in part by the Institut National de la Santé et de la Recherche Médicale (Inserm), Centre National de la Recherche Scientifique (CNRS), University of Lille, Pasteur Institute of Lille. This project was cofounded by the React-EU COVID2I (programme opérationnel FEDER/FSE/IEJ Nord-Pas de Calais) (FT) and the French National Research Agency (Agence Nationale de la Recherche, ANR): AAP générique 2022, ANR-23-CE15-0014-01, GUTSY) (FT). This project was also cofounded by Fundação de Amparo à Pesquisa do Estado de São Paulo (FAPESP, 2018/15313-8) (MARV). PBR received fellowships from FAPESP (2019/14342-7 and 2022/02058-5). VRR received fellowship from FUNCAMP/NIH (1R01DK126969).

### Data availability statement

The sequence datasets (metagenomics) generated in this study are publicly available at https://www.ncbi.nlm.nih.gov/bioproject/PRJNA1121382(https://dataview.ncbi.nlm.nih.gov/object/PRJNA1121382?reviewer=gur7ir3sv1c33coljpo6df56rq). The sequence datasets (transcriptomics) generated in this study are publicly available at https://www.ncbi.nlm.nih.gov/bioproject/1118246. Metabolomics data is deposited to the EMBL-EBI MetaboLights database (DOI: 10.1093/nar/gkad1045, PMID: 37971328) available at https://www.ebi.ac.uk/metabolights/MTBLS10717.

## Acknowledgements

We would like to thank Robin Prath for its technical assistance in the BSL3 laboratory. The authors greatly acknowledge the Lions club from Marcq-en-Baroeul (France) for the purchase of the BSL3 isolator. Biomnigene (Besançon, France) is acknowledged for shotgun sequencing and Fabrice Bouilloux is acknowledged for submitting raw sequence data from the National Center for Biotechnology Information.

## Legend for figures

**Supplementary Figure 1.**
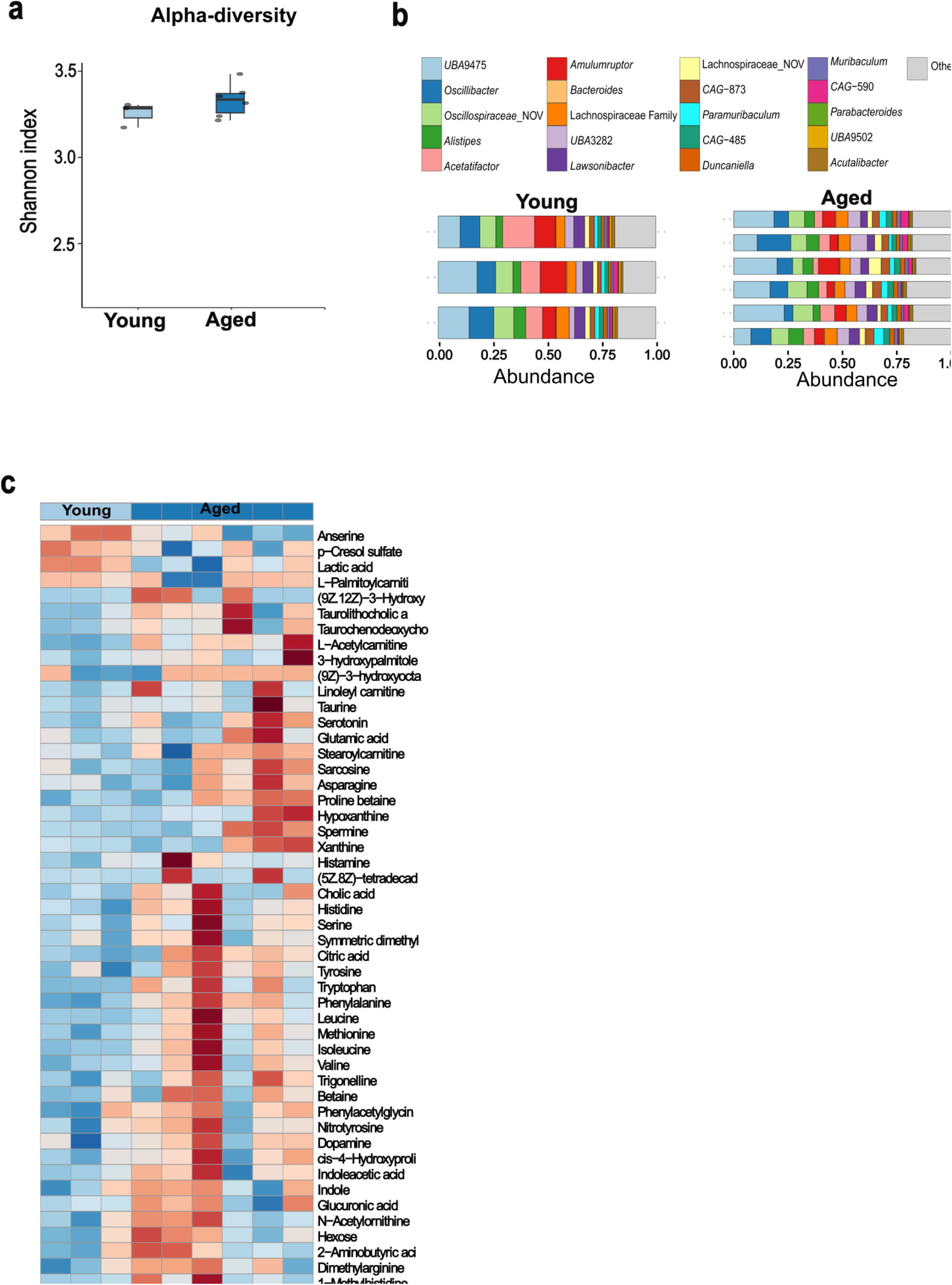
Alpha diversity and taxonomy of gut microbiota and plasma metabolite composition in young and aged hamsters (steady state). **a,** Alpha diversity analysis of gut microbiota communities. Boxplots representing the distribution of the Shannon diversity index for young (light blue) and aged (dark blue) hamsters. **b**, Bar plots showing the distribution of bacterial genera within each sample group. **c,** Heatmap of top 50 metabolites. Each colored cell on the map corresponds to a normalized metabolite concentration, with metabolites name in rows and samples in columns. Young=light blue; Aged=dark blue.

**Supplementary Figure 2.**
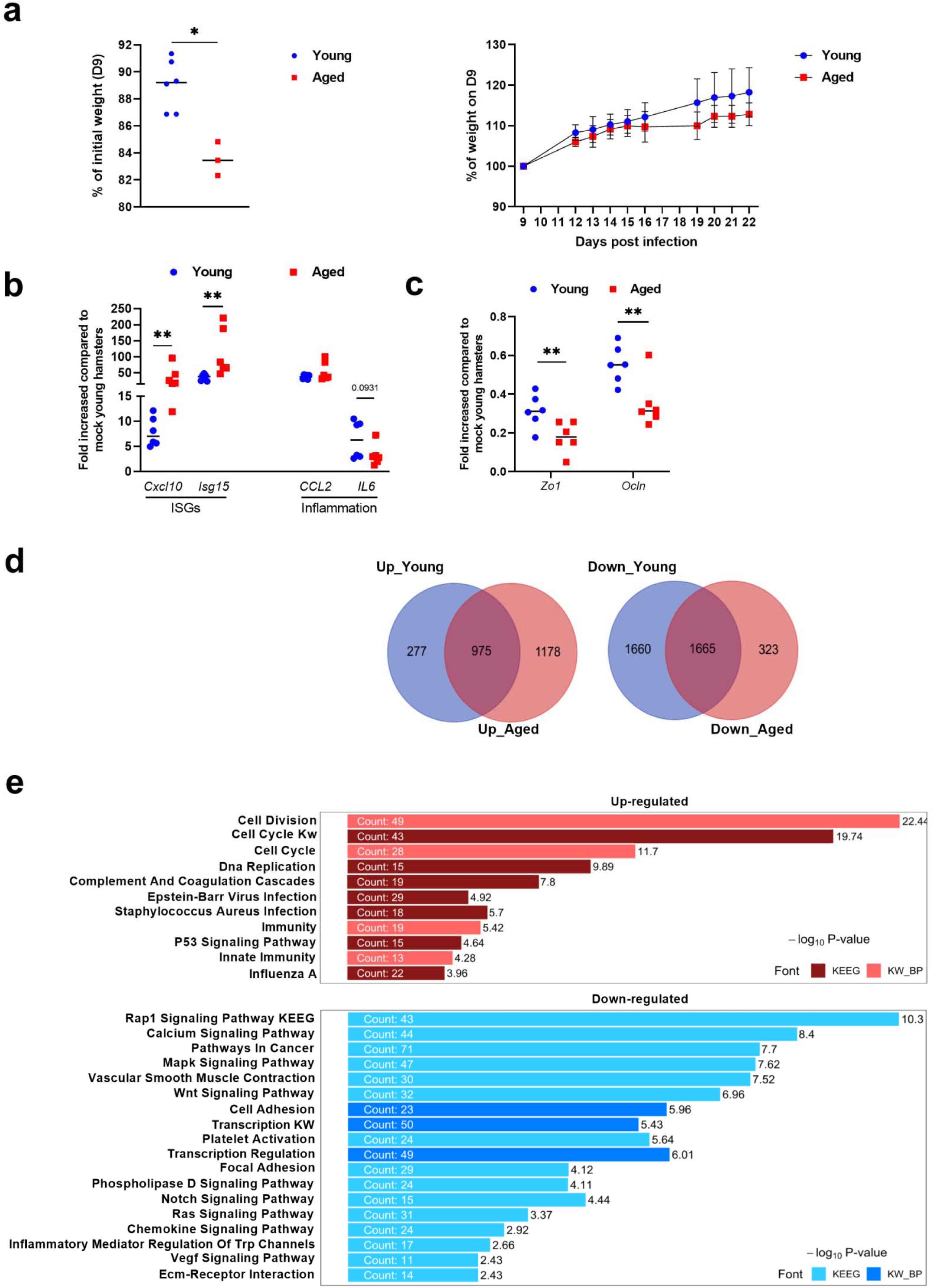
Body weight loss and recovery, expression of fibrotic markers (D22) and lung transcriptomic signatures (D7) of SARS-CoV-2-infected young and aged hamsters. **a**, *Left* panel: Percentage of body weight loss in SARS-CoV-2-infected young and aged hamsters at D8. *Right* panel: Percentage of body weight gain from D8 onwards (6 young hamsters and 3 aged hamsters). **b** and **c**, Lungs from vehicle-treated (mock) and SARS-CoV-2-infeced young and aged were collected on D7. mRNA copy numbers (for ISGs and inflammatory genes in panel **b** and for genes related to barrier functions in panel **c**) were quantified by RT-PCR. The data are expressed as the mean fold change relative to average gene expression in mock-infected young animals (n=6). **d**, Venn diagrams showing the distribution of up-regulated (n=2430) and down-regulated (n=3648) genes at D7. Compared to non-infected age-matched hamsters, 1,252 genes were up-regulated and 3,325 genes were down-regulated in the young group, whereas 2,153 genes were up-regulated and 1,988 were down-regulated in the aged group (*P* < 0.05). **e**, Gene set enrichment analysis from Kyoto Encyclopedia of Genes and Genomes (KEEG) and Keywords Biological Process (KW_BP) (common to both young and aged hamsters) (n=4/group). Regarding common genes shared between young and aged hamsters (40% of upregulated and 45% of downregulated genes, gene ontology (GO) enrichment analysis revealed that upregulated genes were associated with “Immunity”, “Cell cycle/cell division”, and “Complement and coagulation cascades”, while downregulated genes were related to signaling pathways such as “MAPK”, “Chemokine receptor”, “VEGF”, and “Notch signaling”. Significant differences were determined using the Mann Whitney *U* test (**a**, *left* panel, **b** and **c**) (\**P* < 0.05, \*\**P* < 0.01).

**Supplementary Figure 3.**
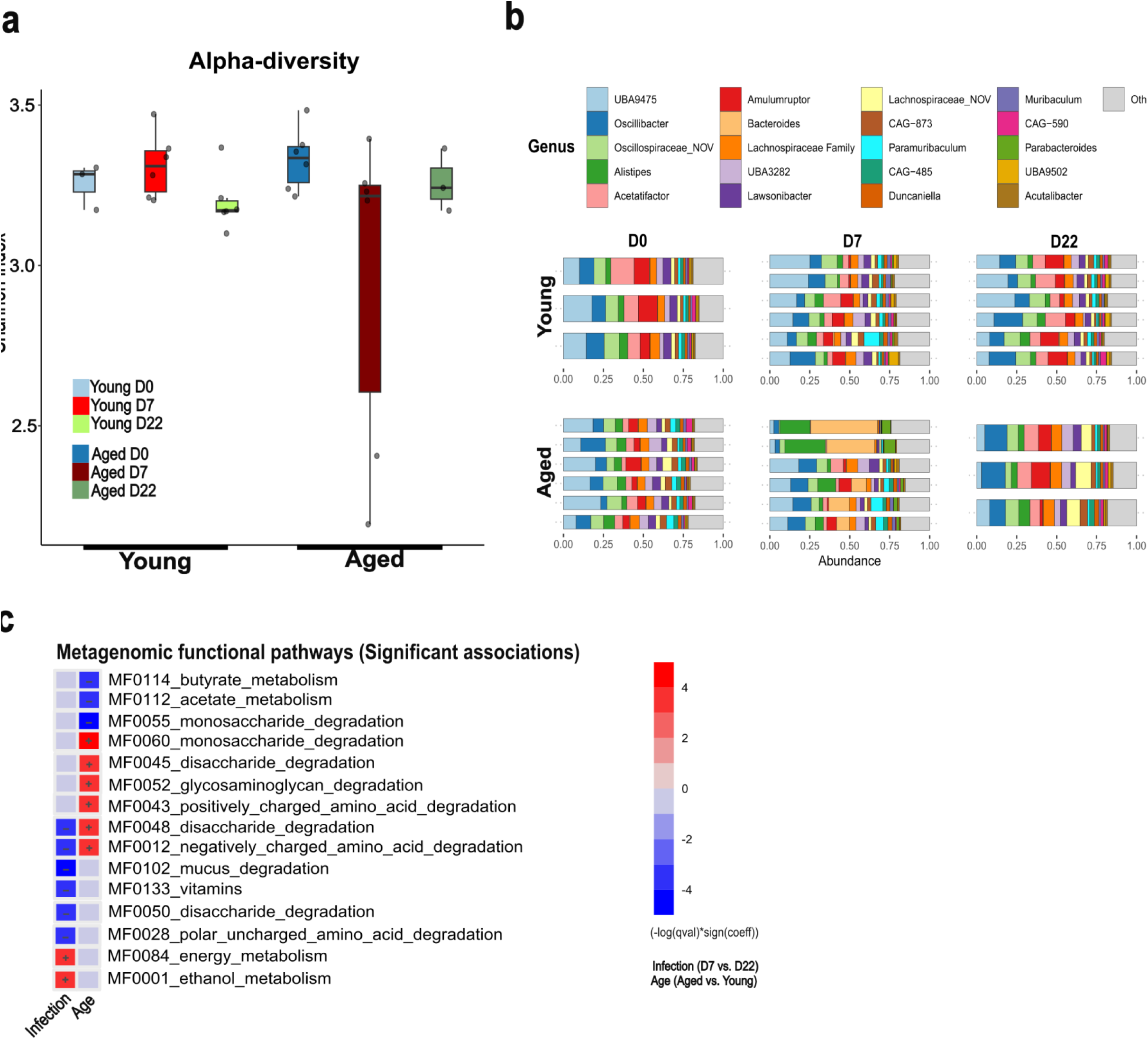
Diversity and functional profile of gut microbiota across different age groups and infection states. **a,** Alpha diversity analysis of gut microbiota communities. Boxplots representing the distribution of the Shannon diversity index for non-infected and infected young and aged hamsters are depicted. **b**, Bar plots showing the distribution of bacterial genera within each sample group. **c**, MaAsLin2 multivariate differential analysis for gut microbiota functional pathways as a heatmap, showing infection and age effects. Positive values (red) indicated enrichment in D22 relative to D7 groups or in aged relative to young groups.

**Supplementary Figure 4.**
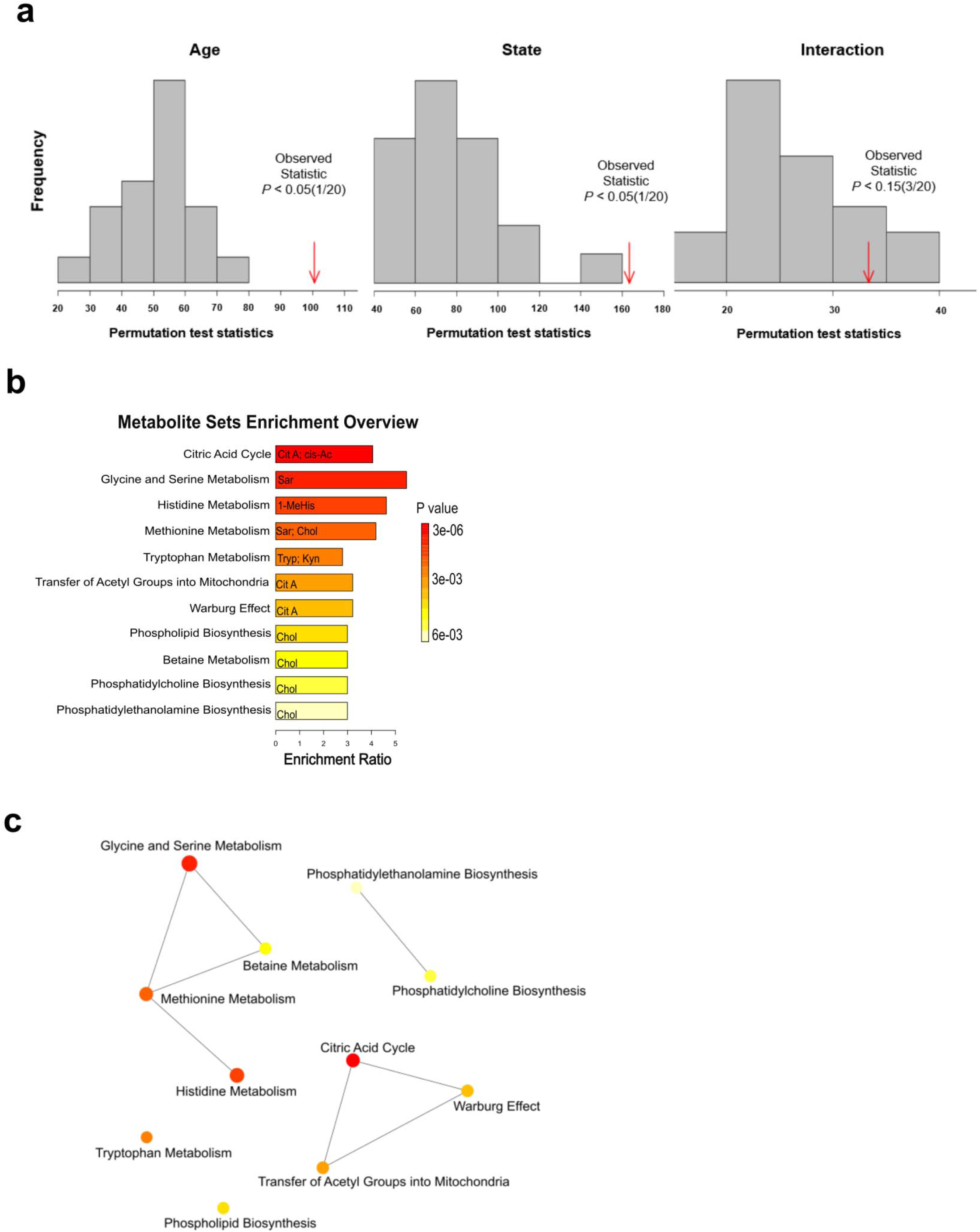
Permutation test and global metabolic network of enrichment pathways from plasma metabolome of infected young and aged hamsters. **a,** Validation by permutation tests based on separation ANOVA-simultaneous component analysis (ASCA). *P* value based on permutation for age, infection state, and their interaction (*P* <0.05). **b**, Pathway analysis based on metabolite sets enrichment performed with the list of significant metabolites (10), identifying the most relevant metabolic pathways via pathway impact and adjusted *P*-value. Figures were drawn using Metaboanalyst software v 5.0. **c,** Global metabolic network of Small Molecular Pathway Database (SMDB) pathways.

**Supplementary Figure 5.**
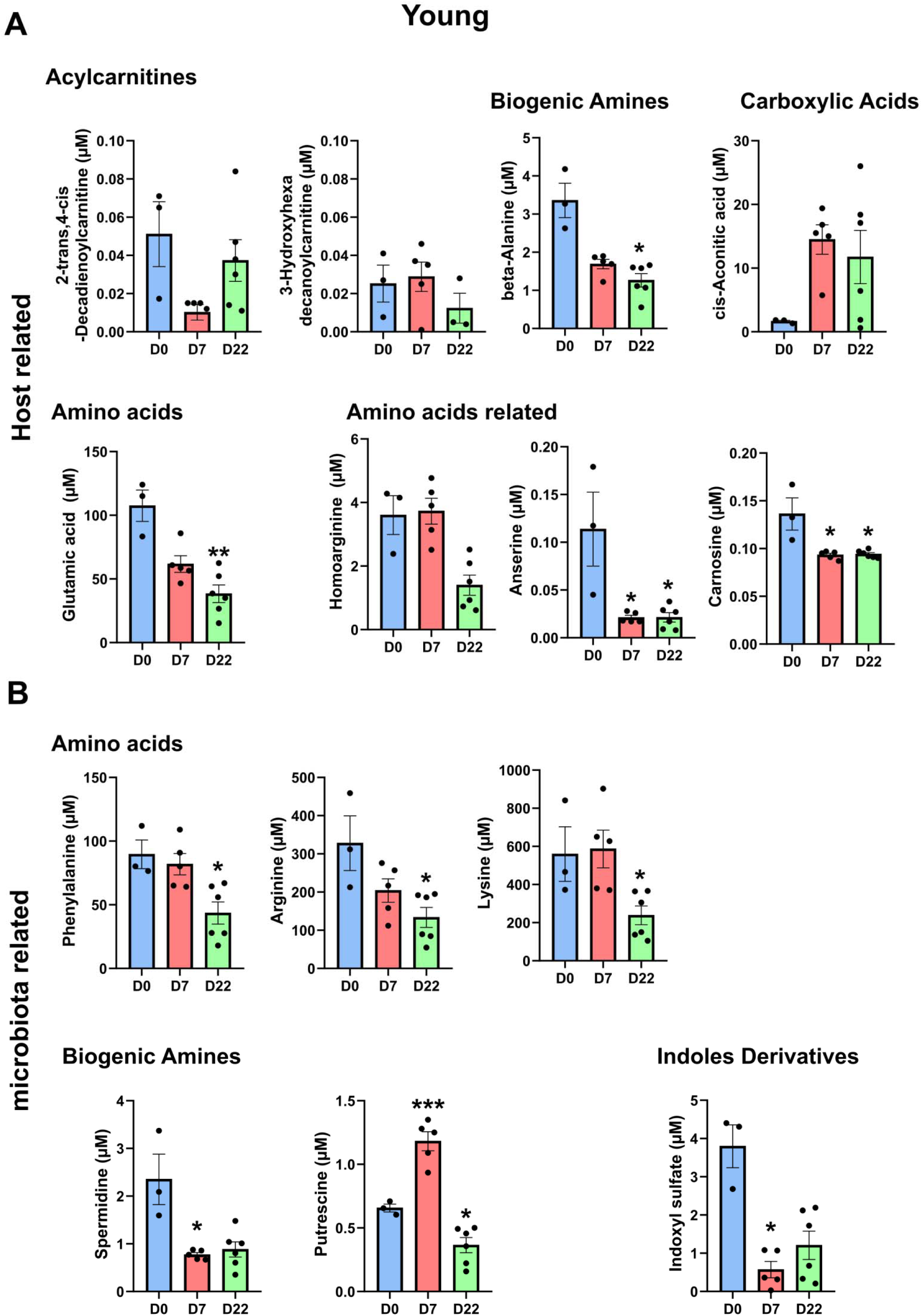
Metabolite concentration shifts in young hamsters post-SARS-CoV-2 infection: insights from network analysis of D7 and D22 post-infection. Plasma concentration of host-related (**a**) and microbiota-related (**b**) metabolites. Data are expressed as mean ± SEM (*n* = 3-5). Significant differences were assessed using the Kruskal-Wallis test, a non-parametric one-way ANOVA, with multiple comparisons (D7 *vs* D0; D22 *vs* D0). Statistical significance is indicated as follows: *P < 0.05; **P < 0.005; ***P<0.001. The bars represent the mean ± SEM (standard error of the mean).

**Supplementary Figure 6.**
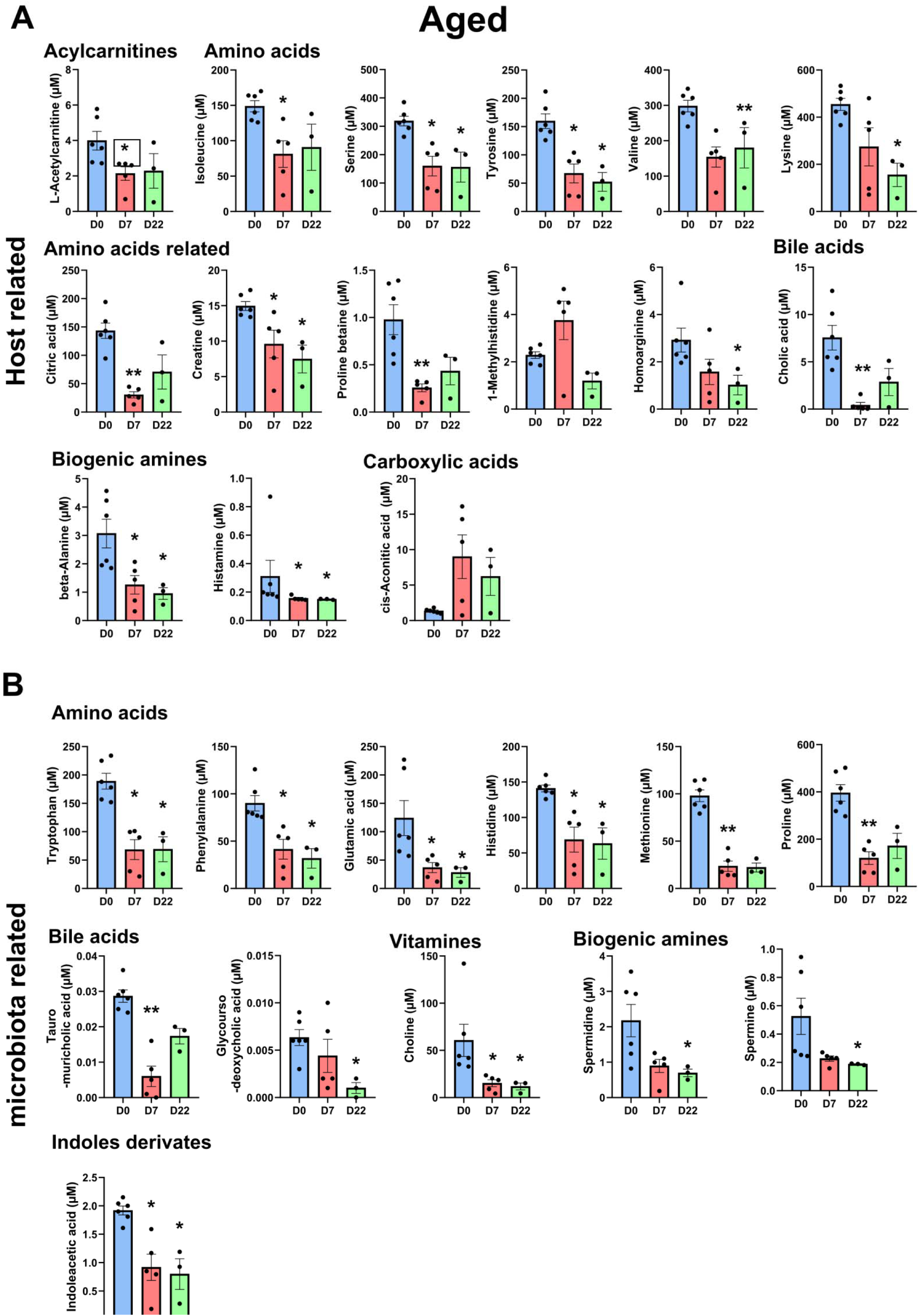
Metabolite concentration shifts in aged hamsters post-SARS-CoV-2 infection: insights from network analysis of D7 and D22 post-infection. Plasma concentration of host-related (**a**) and microbiota-related (**b**) metabolites. Data are expressed as mean ± SEM (*n* = 3-5). Significant differences were assessed using the Kruskal-Wallis test, a non-parametric one-way ANOVA, with multiple comparisons (D7 *vs* D0; D22 *vs* D0). Statistical significance is indicated as follows: *P < 0.05; **P < 0.005; ***P<0.001. The bars represent the mean ± SEM (standard error of the mean).

