## Supplementary material for "Integrative metagenomics and metabolomics reveal age-associated gut microbiota and metabolite alterations in experimental COVID-19": Sup table 4

| *Gapdh* | Forward 5’-CCGTATTGG ACGCCTGGTTA -3’ | *Zo1* | Forward 5’-CTCCTGCCGCTCAAAAGGA -3’ |
| --- | --- | --- | --- |
|  | Reverse 5’-TGAAGTCGCAGGAGACAACC-3’ |  | Reverse 5’-CGCCGGAAGTAGCACCATTA-3’ |
| *Actg1* | Forward 5’-ACAGAGAGAAGATGACGCAGATAATG-3’ | *Ocln* | Forward 5’-GTGGCTTCCACACTTGCTTG-3’ |
|  | Reverse 5’-GCCTGAATGGCCACGTACA-3’ |  | Reverse 5’-GCCACTTCCTGCATAAGGGT-3’ |
| *RdRp* | Forward 5’-GTGARATGGTCATGTGTGGCGG-3’ | *Col1a1* | Forward 5’-CAGCCTACTTCCCCACCTAGC-3’ |
|  | Reverse 5’-CARATGTTAAASACACTATTAGCATA-3’ |  | Reverse 5’-AGGCTCCTTCAAAAGTCCAAGA-3’ |
| *Cxcl10* | Forward 5’-AGGTCAAGAATGCAGGAATGACA-3’ | *Col3a1* | Forward 5’-CCTATGACATCGGTGGTCCG-3’ |
|  | Reverse 5’-AGTGGTGAGGCTACAATGCC-3’ |  | Reverse 5’-CGGTTTTGTTTTGCTGGGGT-3’ |
| *Isg15* | Forward 5’-CTGGTGCCCCTGACTAACTC-3’ | *Ecm1* | Forward 5’-TGGGGACCATATCCAGAGCA-3’ |
|  | Reverse 5’-CTGTCATTCCGCACCAGGAT-3’ |  | Reverse 5’-GGCTTCATCTCTCTCGGCTC-3’ |
| *Ccl2* | Forward 5’-TGCTAACTTGACGCAAGCTCC-3’ | *Mmp2* | Forward 5’-CTCTCGAATCCATGACGGGG-3’ |
|  | Reverse 5’-AAGTTCTTGAGTCTGCGGTGG-3’ |  | Reverse 5’-AACACCAGAGGAAGCCATCG-3’ |
| *Il6* | Forward 5’-CCATGAGGTCTACTCGGCAAA-3’ |  |  |
|  | Reverse 5’-GACCACAGTGAATGTCCACAGATC-3’ |  |  |
